# A Glycosylation-Dependent Checkpoint Restrains Intestinal Intra-Epithelial Lymphocyte Activation

**DOI:** 10.64898/2026.01.02.697332

**Authors:** Liqing Cheng, Niccolò Bianchi, Hadrien Soldati, Ludivine Bersier, Maximilien Evrard, Aurelie Caillon, Amanpreet Singh Chawla, Britt Nakken, Peter Szodoray, Elien De Bousser, Nele Festjens, Tatiana V. Petrova, Sylvain Lemeille, Laura K. Mackay, Nico Callewaert, Mahima Swamy, Simon M.G. Braun, Simone Becattini

## Abstract

Intraepithelial lymphocytes (IELs) are abundant in the intestinal epithelium, where they maintain barrier integrity and provide immune defense. Because of their potent cytotoxic and effector potential, IEL activity must be tightly controlled to prevent tissue damage. However, the mechanisms that calibrate IEL responsiveness remain unclear. Here, we identify glucosaminyl (N-acetyl) transferase 2 (*Gcnt2*) as a key restrainer of both natural and induced gut IEL. Among T cells, *Gcnt2* is uniquely enriched in the intestine and partly dependent on retinoic acid signaling. GCNT2-mediated branched glycosylation marks IELs with signatures of tissue adaptation and reduced TCR responsiveness. Genetic ablation of *Gcnt2* enhanced IEL degranulation, cytokine production, and cytotoxicity upon stimulation, improving bacterial clearance and limiting infection-induced disease, while aggravating the pathological consequences of strong T cell activation. Mechanistically, GCNT2-mediated glycosylation of CD45 reduced its phosphatase activity, thereby dampening TCR signaling and effector responses. Together, these findings reveal GCNT2 as a glycosylation-dependent checkpoint that fine-tunes IEL effector functions, uncovering a novel mechanism by which the intestinal immune system balances responsiveness and tolerance.

**Summary:** GCNT2-mediated I-branching glycosylation of CD45 restrains IEL activation.

## Introduction

The intestinal epithelium contains an abundant population of intraepithelial lymphocytes (IELs) that safeguard barrier integrity and maintain immune homeostasis (Lockhart et al., 2024; McDonald et al., 2018; Van Kaer and Olivares-Villagomez, 2018). IELs comprise two major groups distinguished by developmental origin. Natural IELs (nIELs) arise in the thymus and include both CD8αα⁺TCRγδ⁺ and CD8αα⁺TCRαβ⁺ populations, with the former representing the predominant subset in the small intestine of mice (Cheroutre and Lambolez, 2008). Induced IELs (iIELs), including tissue-resident memory T cells (Trm), derive from conventional peripheral T cells that migrate to the gut epithelium in response to infection, microbial metabolites, or local tissue cues, and include CD8αβ⁺TCRαβ⁺, CD4⁺TCRαβ⁺, and CD4⁺CD8αα⁺TCRαβ⁺ cells (James et al., 2021).

IELs are continuously exposed to antigens from the microbiota, diet, and damaged epithelium, and they integrate diverse activating signals, including TCR ligation (Vandereyken et al., 2020), NK receptor engagement (Lockhart et al., 2024; Yomogida et al., 2024), and cytokines such as IL-18 and IL-15 (James et al., 2021; Montufar-Solis et al., 2007; Okazawa et al., 2004). These cues can rapidly trigger effector functions, supported by the abundant cytotoxic granules characteristic of both γδ and αβ IELs (Cheroutre et al., 2011; Lockhart et al., 2024).

Because of this intrinsic responsiveness, IELs are often described as “activated yet resting”, poised for immediate defense while requiring strict regulatory control to avoid epithelial damage (Shires et al., 2001; Yomogida et al., 2024). Breakdown of this control contributes to immunopathology, including celiac disease and inflammatory bowel disease (Abadie et al., 2020; Dart et al., 2023; Meresse et al., 2006; Xu et al., 2025). Several mechanisms are therefore in place to restrain IEL activation. CD8αα can dampen TCR signaling by engaging thymus leukemia (TL) antigen on epithelial cells and sequestering Lck and LAT (Cheroutre and Lambolez, 2008). nIELs also express attenuated levels of co-stimulatory molecules and altered CD3ζ modules (Cheroutre and Lambolez, 2008; Guy-Grand et al., 1994; Ohteki and MacDonald, 1993; Van Houten et al., 1993; Watt et al., 2023). Additional regulatory pathways connect IEL restraint to metabolic and cytokine inputs, including GLP1R signaling (Wong et al., 2022), and transcriptional programs that limit responsiveness to IL-15 (Yomogida et al., 2024).

Protein glycosylation is an important but underexplored regulatory layer in lymphocytes. Reduced N-glycan branching can enhance TCR signaling and promote autoimmunity and dysregulated glycosylation has been reported in intestinal inflammation (Demetriou et al., 2001; Dias et al., 2018; Smith et al., 2018). *Gcnt2* encodes a β1,6-N-acetylglucosaminyltransferase that generates I-branched poly-N-acetyllactosamine structures, creating multibranched glycan arrays found on both N- and O-linked glycans (Chen et al., 2005; Dimitroff, 2019). In this way, extra initiation points for polyLacNAc-chain elongation are formed, resulting in large multi-branched polyLacNAc arrays, a form of which is present in human erythrocytes and there known as the blood I-antigen (I-Ag) (Bierhuizen et al., 1993). While GCNT2 has been shown to modulate receptor signaling in B cells (Giovannone et al., 2018), its expression pattern, regulation, and function in T cells, particularly in the IEL compartment, have not been defined.

Here, we identify *Gcnt2* as a defining molecular feature of abundant IEL subsets, and a key restrainer of their activation. *Gcnt2* expression marks IELs displaying hallmarks of reduced responsiveness to further stimulation. Loss of *Gcnt2* heightens IEL activation and enhances protection against oral bacterial infection, while being associated with detectable, albeit modest, sequelae of immune activation. Mechanistically, GCNT2-mediated branched glycosylation affects the activity of CD45, a critical tyrosine phosphatase in T cells, dampening TCR signaling. Together, these findings uncover a previously unrecognized glycosylation-dependent checkpoint that governs IELs homeostasis and function.

## RESULTS

### GCNT2-mediated I-branching glycosylation is a defining feature of gut-resident CD8⁺ T cells

Natural and induced IELs share cytotoxic potential and an elevated activation threshold thought to prevent epithelial injury (Lockhart et al., 2024). We reasoned that common mechanisms enforcing this restrained state might be revealed by identifying transcriptional features shared between natural IELs and induced gut Trm cells. To this end, we compared transcriptional programs of endogenous nIELs and iIELs. We generated intestinal tissue-resident memory cells (Trm, iIELs) by transferring naïve CD8⁺ OT-I cells and orally infecting streptomycin-treated mice with *Listeria monocytogenes–OVA* (*Lm*-*OVA*) (Cheng and Becattini, 2024) (**Fig. 1A**), and performed RNA-seq on OT-I cells isolated from spleen and intestinal epithelium at day 30. We integrated this dataset with a published comparison of endogenous TCRβ^+^CD8αα⁺ IELs versus nIEL precursor cells from the thymus (**Fig. 1B**) (Nie et al., 2022). Across both datasets, 327 genes were commonly upregulated in gut-resident CD8⁺ T cells (**Fig. S1A**), including several glycosyltransferases; among these, *Gcnt2* showed the strongest intestinal enrichment (**Fig. 1, C and D**).

**Figure 1.**
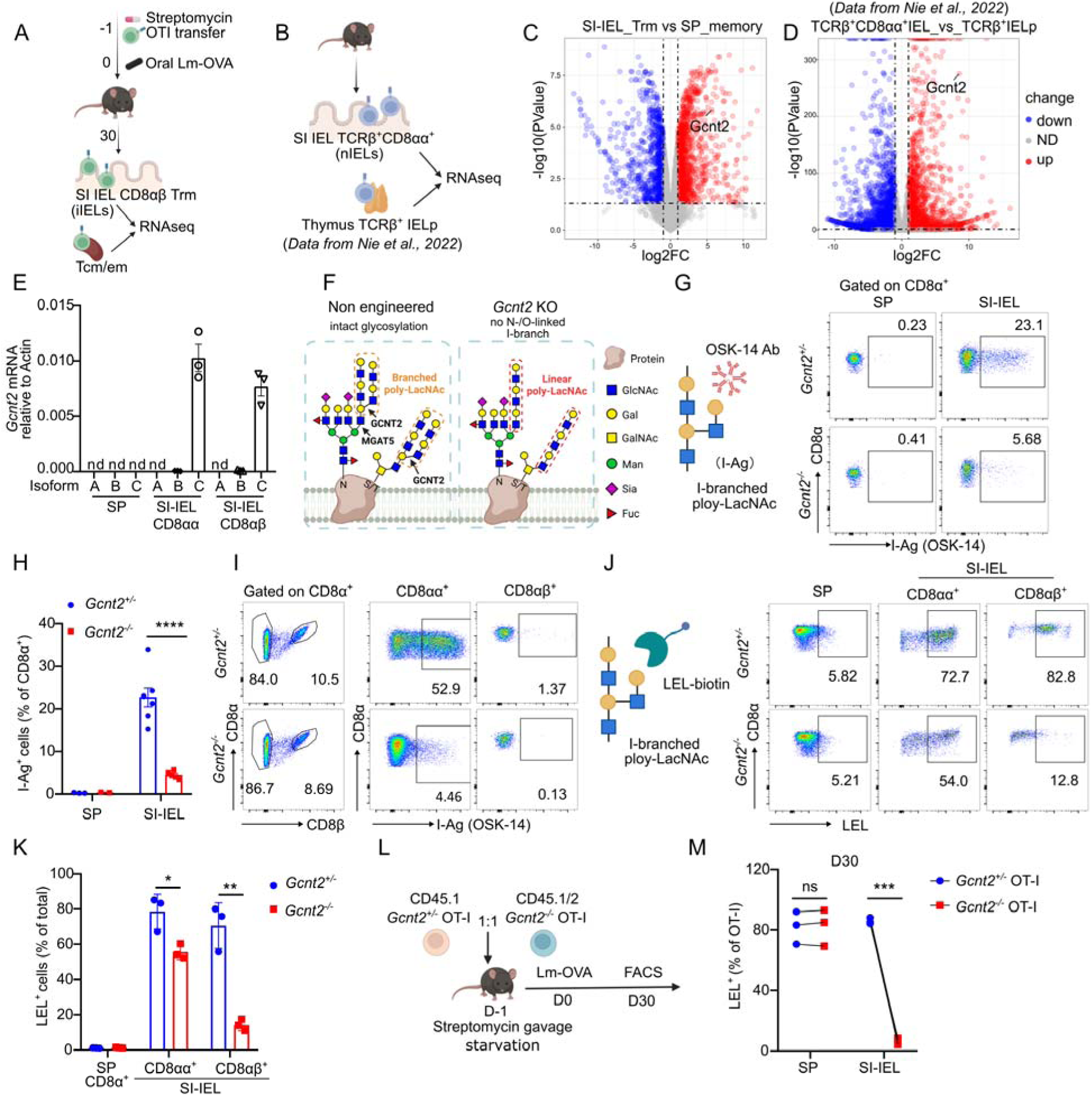
GCNT2-meditated I-branching glycosylation is a defining feature of gut-resident CD8 T cells. (A) Schematic layout of the experiment. OT-I CD8^+^ T cells were transferred into congenically distinct hosts that were orally administered streptomycin and infected 16h later with *Listeria-OVA* (*Lm-OVA*) via gavage. Donor cells from the spleen and small intestine epithelial layer were sorted at day 30 post infection for bulk RNA-seq. (B) Reanalysis of a published RNA-seq dataset (Nie et al., 2022), including endogenous TCRβ^+^CD8αα^+^ IEL from the small intestine and CD4^-^CD8α^-^TCRβ^+^CD5^hi^PD-1^hi^CD122^+^H-2K^b+^CD44^lo^ IEL precursor cells from the thymus. (C) Volcano plot displaying differentially expressed genes between SI-IEL Trm and splenic memory OT-I cells from (A). (D) Volcano plot displaying differentially expressed genes between TCRβ^+^CD8αα^+^ IEL and TCRβ^+^ IEL precursor cells from (B). (E) Isoform-specific RT-qPCR showing expression of *Gcnt2* isoforms in distinct CD8⁺ T cell subsets. (F) Schematic representation of GCNT2-mediated branching glycosylation. (G) Representative FACS plot depicting I-branching glycosylation as detected by OSK-14 antibody staining on CD8α⁺ T cells. (H) Percentage of I-Ag^+^ CD8α^+^ cells in the spleen and SI-IEL from (G). (I) FACS plots depicting OSK-14 staining of CD8αα⁺ and CD8αβ⁺ T cells in SI-IEL. (J) Representative FACS plots showing *Lycopersicon esculentum* lectin (LEL) staining of CD8αα⁺ and CD8αβ⁺ T cells in SI-IEL and spleen. (K) Quantification of LEL staining in the indicated populations from (J). (L) CD45.1 *Gcnt2^+/-^* and CD45.1/2 *Gcnt2^-/-^* OT-I cells were co-transferred into CD45.2 WT recipients, followed by oral *Lm*-OVA infection. At day 30, LEL staining on OT-I cells was analyzed in different organs by FACS. (M) Quantification of LEL⁺ OT-I cells in spleen and SI-IEL from (L). Data in (E) and (K) are one representative experiment of two independent experiments with n=3. Data in (H) are pooled from two independent experiments with n=5-6. Data in (M) are one representative experiment of four independent experiments with n=3. Data are shown as mean ± SEM. Statistical analysis: unpaired t test. *p<0.05; **p<0.01; ***p<0.001; ****p<0.0001; ns, not significant.

Analysis of published scRNA-seq datasets confirmed selective *Gcnt2* expression in small intestinal CD8⁺ T cells relative to mLN (**Fig. S1B**) (Villa et al., 2024), and confinement to the epithelium in Trm across tissues (Crowl et al., 2022; Milner et al., 2020) (**Fig. S1, C and D**), suggesting specific activity regulation in IELs. *Lm*-*OVA*-generated OT-I Trm displayed *Gcnt2* expression across the intestinal tract, highest in SI-epithelium (**Fig. S1E**). *Gcnt2* can be expressed in three distinct alternative spliced isoforms, namely A, B and C, with B being predominant in most tissues (Perez et al., 2021). RT–qPCR showed preferential expression of isoform C in T cells, with markedly higher transcript levels in CD8αα⁺ nIELs and CD8αβ⁺ iIELs than in splenic counterparts (**Fig. 1E**).

GCNT2 generates I-branched polyLacNAc structures via addition of β1,6-linked GlcNAc moieties to internal Gal residues (I-antigen, I-Ag) (**Fig. 1F**). Using OSK-14, a human IgM antibody recognizing I-Ag (1990; Chen et al., 2005; Inaba et al., 2003), we detected no signal on intestinal OT-I Trm, but robust staining on endogenous CD8α⁺ IELs (**Fig. S1F**). This staining was gut-restricted and enriched in CD8α⁺ T cells from SI-IEL, SI-LP, and LI-IEL, but absent in *Gcnt2*^-/-^ mice (**Fig. 1, G and H, and Fig. S1, G-J**). Staining in the large intestinal lamina propria (LI-LP) was lower and appeared to be GCNT2-independent, as no difference was seen between *Gcnt2*^+/-^ and *Gcnt2*^-/-^ mice (**Fig. S1, I and J**). Within the intestinal epithelium, OSK-14 staining was confined to CD8αα⁺ T cells, both TCRαβ⁺ and TCRγδ⁺, and absent from CD8αβ⁺, CD4⁺, and CD4⁺CD8αα⁺ T cells (**Fig. 1I and Fig. S1K**).

Because OSK-14 recognition likely depends on glycan valency, possibly missing structures with reduced branching or elongation, we additionally used *Lycopersicon esculentum* lectin (LEL), which binds polyLacNAc structures regardless of branching topology, to retrieve presence of other GCNT2-mediated glycosylation (Giovannone et al., 2018; Sweeney et al., 2018) (**Fig. 1J**). Indeed, LEL staining revealed reduced glycosylation in both SI-IEL CD8αβ⁺ and CD8αα⁺ T cells from *Gcnt2*^-/-^ mice, albeit with increased background (**Fig. 1K**). CD4⁺ and CD4⁺CD8αα⁺ IELs showed similar LEL staining reduction in *Gcnt2*^-/-^animals (**Fig. S1, L and M**). Following transfer of *Gcnt2*^+/-^ or *Gcnt2*^-/-^ OT-I cells and *Lm*-*OVA* infection, *Gcnt2*^+/-^ OT-I Trm cells recovered from the small intestinal epithelium exhibited stronger LEL staining than their *Gcnt2*^-/-^ counterparts, confirming that GCNT2 also modifies glycans on iIELs (**Fig. 1, L and M**).

Accordingly, *Gcnt2*^+/-^ OT-I Trm showed higher LEL staining than *Gcnt2*^-/-^ cells at multiple gut sites (**Fig. S1, N and O**). A summary of OSK-14 versus LEL staining across IEL subsets is shown in **Fig. S1P**. Furthermore, *Gcnt2*^-/-^ IELs displayed enhanced binding of Galectin-9 and *Phaseolus vulgaris Leucoagglutinin* (PHA-L) (**Fig. S1, Q and R**), consistent with previous report that GCNT2-mediated I-branching glycosylation inhibits Galectin-9 and PHA-L binding to cells (Giovannone et al., 2018). In addition to CD8⁺ T cells, GCNT2-dependent polyLacNAc staining was also detected in Th17 cells, ILC3s, γδ T cells, and Tregs in the intestinal lamina propria, while their cellular number remained unaffected in *Gcnt2*^-/-^ mice (**Fig. S2, A-D**). Collectively, these findings indicate that GCNT2 decorates nIELs, iIELs, and additional intestinal immune subsets with branched polyLacNAc structures that may vary in density and/or conformation.

### Intestinal retinoic acid promotes *Gcnt2* expression and branched glycosylation upon gut seeding

Because *Gcnt2* expression and I-branching were specific to intestinal cells and not detectable in lymphoid tissues, we hypothesized that signals encountered upon gut entry induced *Gcnt2*. Consistent with this, OSK-14 and LEL staining were absent in the thymus (**Fig. 2, A and B**). Transfer of IEL precursors (including both CD4^-^CD8^-^TCRβ^+^ and CD4^-^CD8^-^TCRγδ^+^ thymocytes) into *Rag2*^-/-^ hosts (Cheroutre et al., 2011), established that I-Ag appeared only in intestinal IELs derived from *Gcnt2*-sufficient donors (**Fig. 2, C-E**), indicating that gut-specific signals induce *Gcnt2*. In line with this hypothesis, a published time course scRNA-seq dataset from a P14 cell transfer and LCMV infection model showed early induction of *Gcnt2* in gut Trm by day 4 post-infection (**Fig. S3A**). Consistently, in our model, transferred OT-I cells recovered from the intestine exhibited increased LEL staining by day 9 after *Lm*-*OVA* infection (**Fig. S3, B and C**).

**Figure 2.**
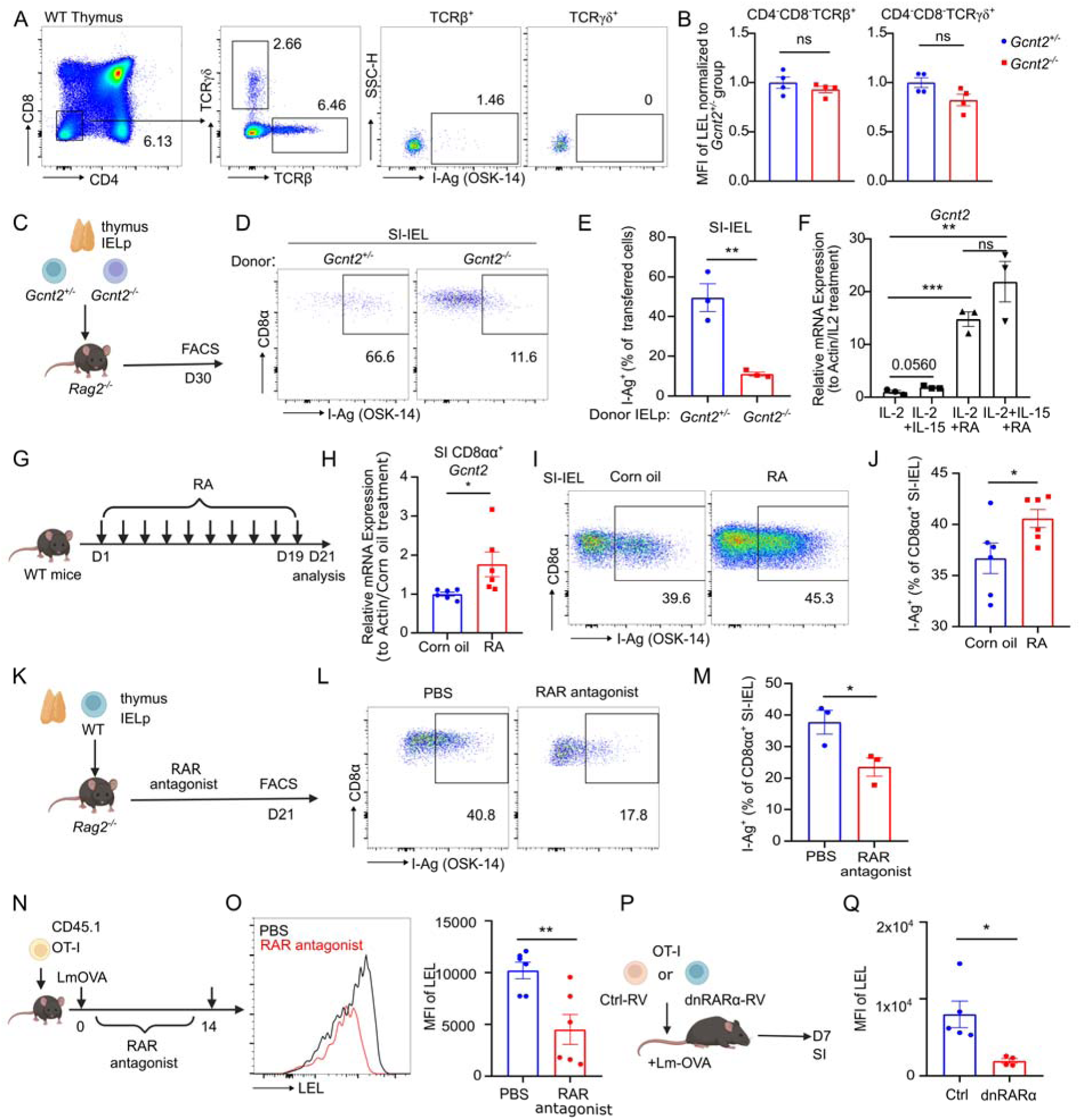
Retinoic acid promotes *Gcnt2* expression and I-branching glycosylation in SI-IEL CD8 T cells. (A) FACS plot showing I-branching glycosylation detected by OSK-14 antibody staining on TCRβ and TCRγδ precursors in the thymus. (B) Normalized MFI of LEL staining on thymic TCRβ and TCRγδ precursors from *Gcnt2*^+/-^ and *Gcnt2*^-/-^ mice. (C) Schematic of IEL precursor transfer into *Rag2^-/-^* recipients for nIEL reconstitution. (D) FACS plot showing I-branching glycosylation detected by OSK-14 antibody staining on transferred donor-derived IELs, as described in (C). (E) Quantification of I-Ag⁺ donor-derived cells in SI-IEL from (D). (F) *Gcnt2* induction by IL-2, IL-15 and RA treatment *in vitro* analyzed by RT-qPCR after long-term (6 days). (G) WT mice were gavaged with either corn oil or RA (200 µg) in corn oil every other day for 20 days. (H) *Gcnt2* mRNA levels in sorted SI-IEL CD8αα⁺ T cells from (G), analyzed by RT-qPCR. (I) I-branching glycosylation staining using OSK-14 antibody was detected by FACS from vehicle or RA treated mice from (G). (J) Percentage of I-Ag^+^ SI-IEL CD8αα^+^ T cells were quantified from (I). (K) WT IEL precursors were transferred to *Rag2^-/-^* recipients, followed by intraperitoneal injection of vehicle or RAR antagonist (1 mg/kg) every other day for 20 days. (L) I-branching glycosylation staining using OSK-14 antibody was detected by FACS from vehicle or RAR antagonist treated mice in (K). (M) Percentages of I-Ag^+^ SI-IEL CD8αα^+^ T cells quantified from (L). (N) CD45.1 WT OT-I cells were intravenously transferred into CD45.2 WT recipients, followed by oral infection with *Lm-OVA* and treatment with vehicle or RAR antagonist (1 mg/kg) every other day for 14 days. (O) Representative FACS plots and MFI of LEL staining on transferred OT-I cells in SI-IEL from vehicle- or RAR antagonist–treated mice in (N). (P)-(Q) CD45.1 WT OT-I cells transduced with either empty vector or dnRARα were transferred to CD45.2 WT recipients infected with *Lm*-*OVA* (P). At day 7 post-infection, SI-IEL OT-I cells were isolated and MFI of LEL staining was quantified (Q). Data in (B), (H), (J), (O) and (Q) are pooled from two independent experiments with n=4-6. Data in (E) and (M) represent one of two independent experiments with n=3. Data in (F) are one representative of two independent experiments. Data are shown as mean ± SEM. Statistical analysis: unpaired t test. *p<0.05; **p<0.01; ***p<0.001; ns, not significant.

We then asked which gut-specific signals may be involved in regulation of *Gcnt2* expression and GCNT2-mediated glycosylation. Commensal microbes of the intestine can impact IEL numbers and features (Alonso et al., 2025; Cervantes-Barragan et al., 2017; Jia et al., 2022; Song et al., 2023), however germ-free (GF) or antibiotic-treated mice did not display alterations in I-branching staining (**Fig. S3, D and E**). We therefore turned our attention to gut-enriched soluble factors. TGF-β, critical for IEL development (Han et al., 2023; Konkel et al., 2011; Mackay et al., 2015; Zhang and Bevan, 2013), did not affect *Gcnt2* expression in CD8^+^ T cells *in vitro* (**Fig. S3, F-I**). Retinoic acid (RA) is an alternative factor that can imprint features of tissue residency in T cells and is known to act on IELs (Obers et al., 2024; Qiu et al., 2023; Reis et al., 2014; Sujino et al., 2016). Mining a recently published dataset we found that RA, but not TGF-β treatment, increased *Gcnt2* mRNA in IL-2 and IL-15-cultured CD8^+^ T cells *in vitro* (**Fig. S3J**) (Obers et al., 2024). Consistently, RA alone or combined with IL-15 strongly induced *Gcnt2* in activated OT-I cells (**Fig. 2F**).

*In vivo*, RA supplementation increased *Gcnt2* mRNA levels and glycosylation in SI-IEL CD8αα⁺ T cells (**Fig. 2, G-J**). Conversely, RA receptor (RAR) antagonist treatment of *Rag2*^-/-^ recipients reconstituted with IEL precursors reduced total IEL numbers and I-branching glycosylation, in both small (**Fig. 2, K-M**) and large intestine (**Fig. S3, K-M**), confirming a role for RA in this process. Short-term RA signaling blockade modestly reduced *Gcnt2* mRNA without affecting staining (**Fig. S3, N-Q**), suggesting either low enzyme turnover, or that *Gcnt2* transcripts are present in excess, so that a twofold decrease in expression is insufficient to alter glycosylation. Similarly, in OT-I-transferred, *Lm*-*OVA* orally infected mice, RAR antagonist administration reduced both Trm accumulation and LEL staining (**Fig. 2, N and O, and Fig. S3, R and S**). Importantly, OT-I cells expressing a dominant-negative RARα (dn- RARα) (Obers et al., 2024), displayed reduced LEL staining early after infection (**Fig. 2, P and Q**), suggesting intrinsic activity of RA on IELs. We conclude that RA, possibly among other gut-specific signals, promotes the expression of *Gcnt2* and polyLacNAc branching in IELs.

### I-antigen marks an intestinal IEL subset with unique molecular features

We next examined the role of *Gcnt2* in IELs. Firstly, conventional T cell development was normal in *Gcnt2*^-/-^ mice (**Fig. S4, A-C**), and numbers of both TCRαβ^+^ and TCRγδ^+^ thymic IEL precursors were comparable (**Fig. S4, D-F**). Interestingly, *Gcnt2*^-/-^ mice showed increased SI-IEL numbers, in particular for CD8αα^+^TCRγδ^+^ subsets (**Fig. S4, G-I**), suggesting a compartmentalized impact of GCNT2 activity. IEL localization along the villus remained unchanged between groups (**Fig. S4J**). Further analysis displayed no differences in TCR Vγ usage or maturation marker expression, such as CD44, between *Gcnt2*^+/-^ and *Gcnt2*^-/-^ mice (**Fig. S4K and L**).

Bulk RNA-seq of total CD8αα⁺ nIELs from *Gcnt2*^+/-^ and *Gcnt2*^-/-^ mice revealed no major transcriptional differences (**Fig. S5A**), suggesting that *Gcnt2* expression does not impact IEL transcriptional profile at steady state. However, reanalysis of two independent scRNA-seq datasets revealed that *Gcnt2*-high CD8α⁺ IELs expressed high *Gzma/b* and low *Tcf7* (**Fig. 3A and Fig. S5, B and C**) (Hioki et al., 2025; Yakou et al., 2023), suggesting that *Gcnt2* expression may identify a functionally distinct IEL subset rather than directly controlling its transcriptional state. Consistent with this interpretation, OSK-14-sorted I-Ag⁺ CD4⁻CD8αα⁺ IELs expressed higher levels of *Gcnt2* and *Gzma/b*, as confirmed at both the transcript and protein levels (**Fig. 3, B-D**), together with lower *Tcf7* expression (**Fig. 3B**). Conversely, CD4⁻CD8αα⁺ IELs from *Gcnt2*^⁻/⁻^ mice expressed granzyme levels comparable to controls (**Fig. S5, D and E**) and nIELs from *Gzma/b* double-knockout mice displayed normal *Gcnt2* transcript levels (**Fig. S5F**), indicating that *Gcnt2* and *Gzma/b* are co-expressed but do not reciprocally regulate each other. Thus, while *Gcnt2* is not required to maintain the steady-state transcriptional profile of CD8αα⁺ IELs, its expression marks a seemingly cytotoxic, Tcf7-low IEL subset that is preserved in *Gcnt2*-deficient mice but can no longer be identified through *Gcnt2*-dependent I-Ag staining.

**Figure 3.**
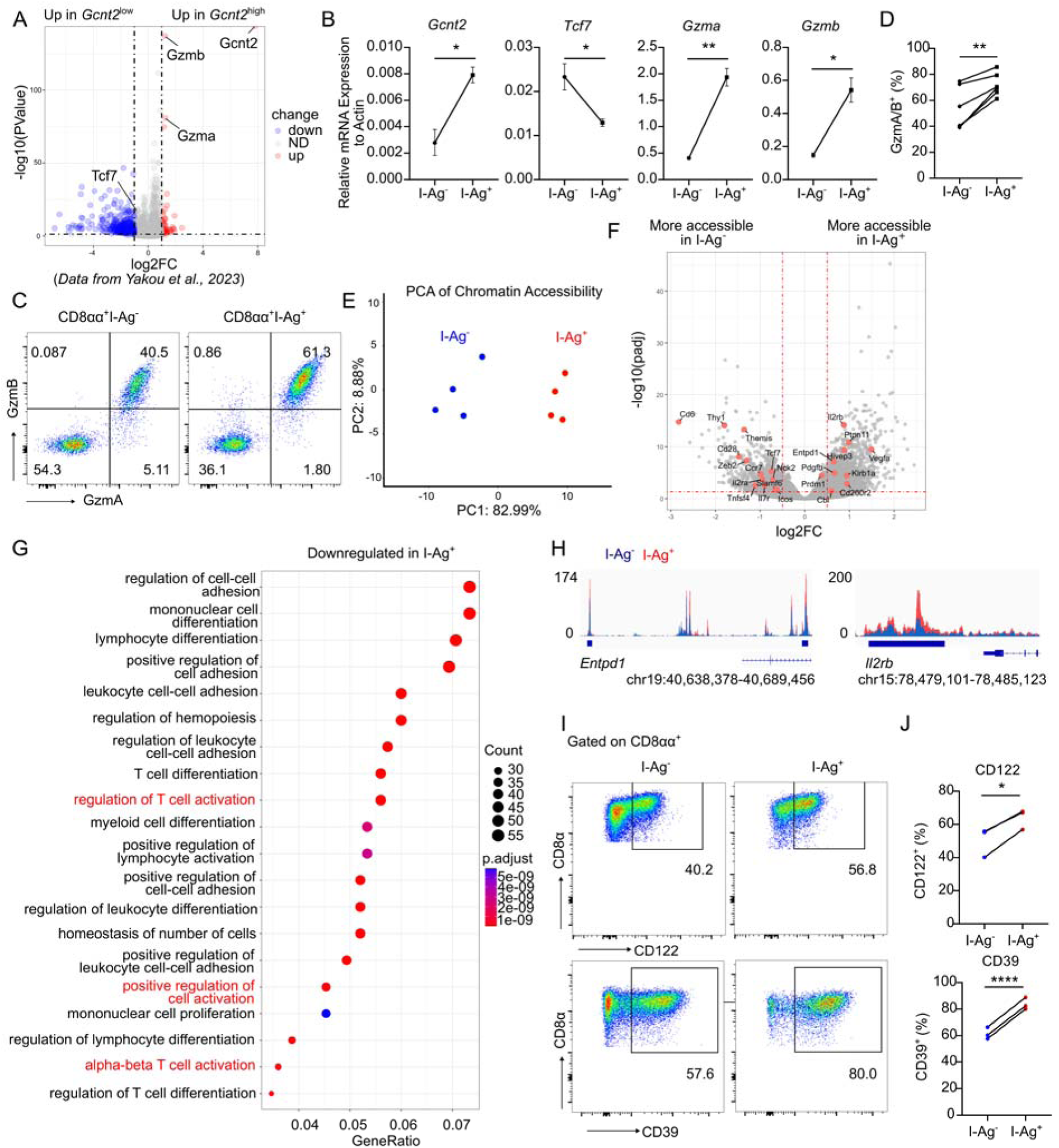
I-branching glycosylation marks a subpopulation of IELs with unique molecular features. (A) Reanalysis of a published CD45^+^ IEL scRNA-seq dataset (Yakou et al., 2023), in which cells were stratified into *Gcnt2^hi^* and *Gcnt2^low^*. Differentially expressed genes between these two populations are shown. (B) I-Ag^+^ and I-Ag^-^ SI-IEL CD4^-^CD8αα^+^ T cells were sorted and RT-qPCR was performed to detect expression of *Gzma*, *Gamb*, *Tcf7* and *Gcnt2*. (C) Representative FACS plot depicting GzmA/B expression in I-Ag^+^ versus I-Ag^-^ SI-IEL CD4^-^CD8αα^+^ T cells, based on OSK-14 staining. (D) Compiled data and statistical analysis from (C). (E) ATAC-seq was performed on OSK-14-sorted I-Ag^+^ and I-Ag^-^ SI-IEL CD4^-^CD8αα^+^ T cells from WT mice, and principal component analysis (PCA) of ATAC-seq samples is displayed. (F) Volcano plot depicting genes that display differential chromatin accessibility between I-Ag^+^ and I-Ag^-^SI-IEL CD4^-^CD8αα^+^ T cells (adjusted p-value < 0.05 and | fold change| > 1.5, n=4), highlighted genes are discussed in the text. (G) Gene set enrichment analysis of pathways downregulated in I-Ag^+^ cells, based on ATAC-seq result. (H) Genome browser tracks of chromatin accessibility data at *Cd39* and *Cd122* genomic loci. (I) and (J) Representative FACS plot (I) and quantification (J) of CD39 and CD122 expression in I-Ag⁺ and I-Ag⁻ SI-IEL CD4^-^CD8αα⁺ T cells. Data in (B), (D) and (J) represent one of two independent experiments with n=3-4. Data are shown as mean ± SEM. Statistical analysis: unpaired t test. *p<0.05; **p<0.01; ****p<0.0001.

To further define the molecular features of this I-Ag⁺ subset, we performed ATAC-seq on sorted I-Ag⁺ and I-Ag⁻ CD4⁻CD8αα⁺ IELs from WT mice, thereby comparing chromatin accessibility between these populations independently of *Gcnt2* deficiency. This analysis revealed clear differences in chromatin accessibility between I-Ag⁺ and I-Ag⁻ IELs (**Fig. 3, E and F**), with most differentially accessible regions mapping to enhancer elements (**Fig. S5G**). The *Gcnt2* locus was modestly but significantly more accessible in I-Ag⁺ IELs, especially at the first exon of isoform C (**Fig. S5H**). Although *Gzma* and *Gzmb* showed no accessibility differences, loci associated with stemness and central-memory features (*Ccr7*, *Il7r*, *Tcf7*) were markedly less accessible (**Fig. 3F and Fig. S5I**). Co-stimulatory and activation-associated genes (*Il2ra*, *Cd6*, *Icos*, *Cd28*, *Slamf6*, *Tnfsf4*) and positive regulators of TCR signaling (*Themis*, *Nck2*) also displayed reduced accessibility, whereas inhibitory receptors (*Cd200r2*, *Klrb1a*) and negative regulators (*Cbl*, *Ptpn11*) were more accessible (**Fig. 3F and Fig. S5J**). Pathway analysis confirmed globally diminished accessibility at TCR signaling loci in I-Ag⁺ IELs (**Fig. 3G**). In contrast, regions encoding *Entpd1* (CD39) and *Il2rb* (CD122) were more accessible in this population (**Fig. 3H**), with correspondingly higher CD39 and CD122 protein levels (**Fig. 3, I and J**). CD39, classically linked to immunoregulation (Borsellino et al., 2007; Timperi and Barnaba, 2021), has recently been associated with IL-15- (and CD122)-driven maturation of IELs exhibiting restrained cytokine production (Alonso et al., 2025). The combination of increased *Il2rb* and reduced *Il2ra* accessibility further suggests a shift toward IL-15 rather than IL-2 responsiveness, a hallmark of innate-like or tissue-adapted IEL states. Together, these data delineate I-Ag⁺ CD8αα⁺ IELs as a transcriptionally and epigenetically distinct subset marked by attenuated activation and co-stimulatory programs, strengthened immunoregulatory and tissue-adaptation features, and unexpectedly high granzyme content, highlighting a specialized state with limited classical responsiveness yet preserved context-dependent cytotoxic potential.

### *Gcnt2* expression restrains IEL activation upon TCR stimulation

ATAC-seq result indicates a potential link between *Gcnt2* expression and reduced IEL responsiveness, which is consistent with prior work implicating polyLacNAc branching in tuning TCR signaling (Demetriou et al., 2001; Dias et al., 2018; Smith et al., 2018). To test this directly, we stimulated sorted CD4^-^CD8αα⁺ IELs from WT mice with anti-CD3. Despite their higher GZMA/B content, I-Ag⁺ cells showed markedly reduced degranulation and cytokine production compared to I-Ag⁻ counterparts (**Fig. 4, A and B**). To determine whether *Gcnt2* functions as a regulator rather than a passive marker of reduced responsiveness, we analyzed CD4^-^CD8αα⁺ IELs from *Gcnt2*^-/-^ mice. CD4^-^CD8αα⁺ IELs lacking *Gcnt2* mounted significantly stronger degranulation and cytokine responses upon anti-CD3 stimulation (**Fig. 4, C and D**), and displayed enhanced cytotoxicity toward YAC1 and A20 lymphoma targets (**Fig. 4E and Fig. S6, A-C**), indicating that GCNT2 plays a direct role in restraining IEL activation.

**Figure 4.**
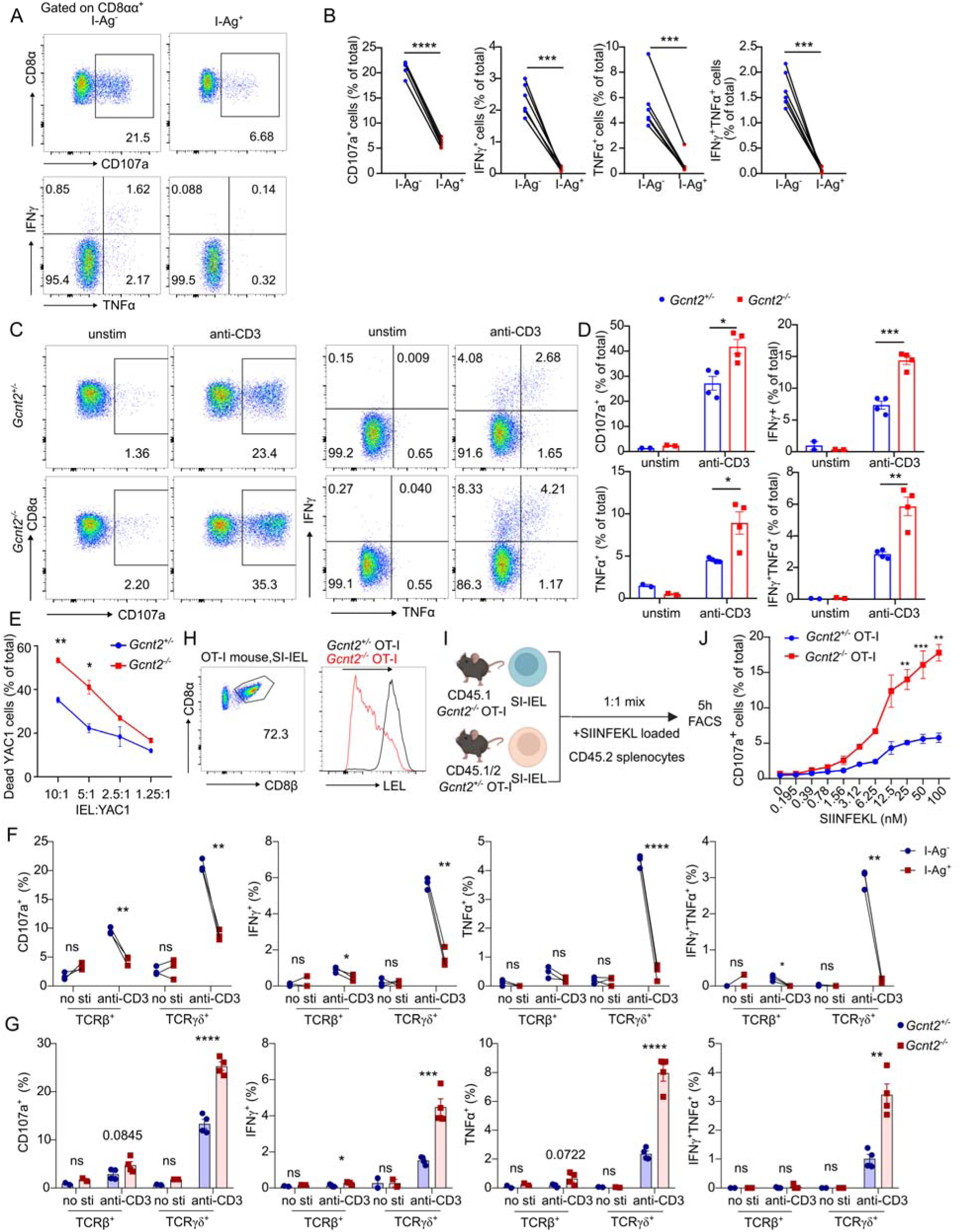
*Gcnt2* expression restrains IELs activation. (A) Sorted I-Ag^-^ or I-Ag^+^ SI-IEL CD4^-^CD8αα^+^ T cells from WT mice were stimulated by plate-bound anti-CD3 (5 µg/ml) in presence of CD107a antibody and Golgi inhibitor for six hours. Degranulation and cytokine production were assessed by FACS. (B) Quantification of data shown in (A). (C-D) Sorted SI-IEL CD4^-^CD8αα^+^ T cells from *Gcnt2*^+/-^ and *Gcnt2*^-/-^ mice were stimulated as in (A). Degranulation and cytokine production were analyzed by FACS and quantified (D). (E) Cytotoxicity assay showing percentage of dead YAC1 target cells after co-culture with anti-CD3-activated IELs at the indicated effector-to-target ratios. (F) Degranulation and cytokine production by I-Ag^+^ vs I-Ag^-^ subsets among CD4^-^CD8αα^+^TCRβ^+^ or CD4^-^CD8αα^+^TCRγδ^+^ SI-IEL cells from *Gcnt2*^+/-^ mice. (G) Degranulation and cytokine production by CD4^-^CD8αα^+^TCRβ^+^ or CD4^-^CD8αα^+^TCRγδ^+^ SI-IEL cells from *Gcnt2^+/-^* and *Gcnt2^-/-^* mice. (H) Representative FACS plots depicting CD8⁺ T cell subset composition in SI-IELs from OT-I mice and LEL staining on SI-IEL CD8αβ⁺ T cells from *Gcnt2*^+/-^ and *Gcnt2*^-/-^ OT-I mice. (I) Schematic representation of degranulation assay in which congenically distinct *Gcnt2^+/-^* and *Gcnt2^-/-^*OT-I SI-IELs were co-cultured with SIINFEKL-loaded splenocytes in the presence of CD107a antibody and Golgi inhibitor. (J) Quantification of data from (I). Data in (B) are pooled from two independent experiments with n=6. Data in (D) represent one of three independent experiments. Data in (E), (F), (G) and (J) represent one of two independent experiments. Data are shown as mean ± SEM. Statistical analysis: unpaired t test. *p<0.05; **p<0.01; ***p<0.001; ****p<0.0001; ns, not significant.

The above conclusions were drawn for total CD4^-^CD8αα⁺ IELs, including both TCRαβ^+^ and TCRγδ^+^ cells, which are functionally distinct (Lockhart et al., 2024). We therefore distinguished the effects of *ex vivo* anti-CD3 stimulation on TCRαβ^+^ and TCRγδ^+^ cells, by comparing both I-Ag⁺ and I-Ag⁻ CD4^-^CD8αα⁺ IELs within WT mice, as well as *Gcnt2*^+/-^ and *Gcnt2*^-/-^ IELs. These experiments confirmed that GCNT2-mediated glycosylation exerts an inhibitory effect on T cells, and uncovered that this effect is prominent in CD4^-^CD8αα⁺TCRγδ^+^ IELs, which also respond more vigorously to the stimulation, and less pronounced in CD4^-^CD8αα⁺TCRαβ^+^ IELs (**Fig. 4, F and G**). This is consistent with the known regulatory rather than inflammatory potential of CD4^-^CD8αα⁺TCRαβ^+^ cells (Klose et al., 2014; Nie et al., 2022; Poussier et al., 2002).

Because anti-CD3 provides a supraphysiological trigger, and most CD4^-^CD8αα⁺ IELs express γδ TCRs responsive to non-classical cues whose nature remains elusive (Vandereyken et al., 2020), we next adopted a model of cognate-antigen stimulation. Using OT-I mice, in which IELs predominantly consist of CD8αβ⁺TCRαβ⁺ cells capable of responding to SIINFEKL (Huang et al., 2011; Leishman et al., 2001), we found that *Gcnt2*^-/-^ OT-I IELs exhibited reduced polyLacNAc density and increased degranulation following cognate peptide stimulation (**Fig. 4, H-J and Fig. S6D**). Together, these data indicate that *Gcnt2* marks abundant yet functionally distinct IEL subsets and that its expression actively restrains effector functions, including degranulation and inflammatory cytokine production, upon TCR triggering.

### *Gcnt2* deficiency improves protection from intestinal pathogens

Natural and induced IELs contribute critically to early defense against intestinal pathogens, including *Salmonella Typhimurium* (STm), by limiting bacterial expansion, preventing epithelial translocation, and promoting tissue repair (Chawla et al., 2025; Dalton et al., 2006; Ebert and Roberts, 1996; Edelblum et al., 2015; Hoytema van Konijnenburg et al., 2017; Ismail et al., 2011; Jia et al., 2022; Li et al., 2012; Sheridan and Lefrancois, 2010; Swamy et al., 2015). Given the heightened effector responsiveness of *Gcnt2*^-/-^ IELs *ex vivo*, we asked whether this translated into improved host protection. Following oral STm infection, *Gcnt2*^-/-^ mice showed reduced weight loss and lower fecal pathogen burden compared with controls, although survival was unchanged (**Fig. 5, A-C**). To determine whether this effect extended to other enteric pathogens, we used an oral *Listeria monocytogenes* (*Lm*) infection model. Here, *Gcnt2*^-/-^mice again displayed enhanced resistance, with reduced weight loss, diminished fecal burden, and improved survival (**Fig. 5, D-F**). Depletion of circulating CD8⁺ T cells using anti-CD8α antibody, an approach that spares IELs while targeting peripheral compartments (**Fig. S7, A-D**), did not alter these outcomes, indicating that early protection may be mediated largely by resident intestinal T cells.

**Figure 5.**
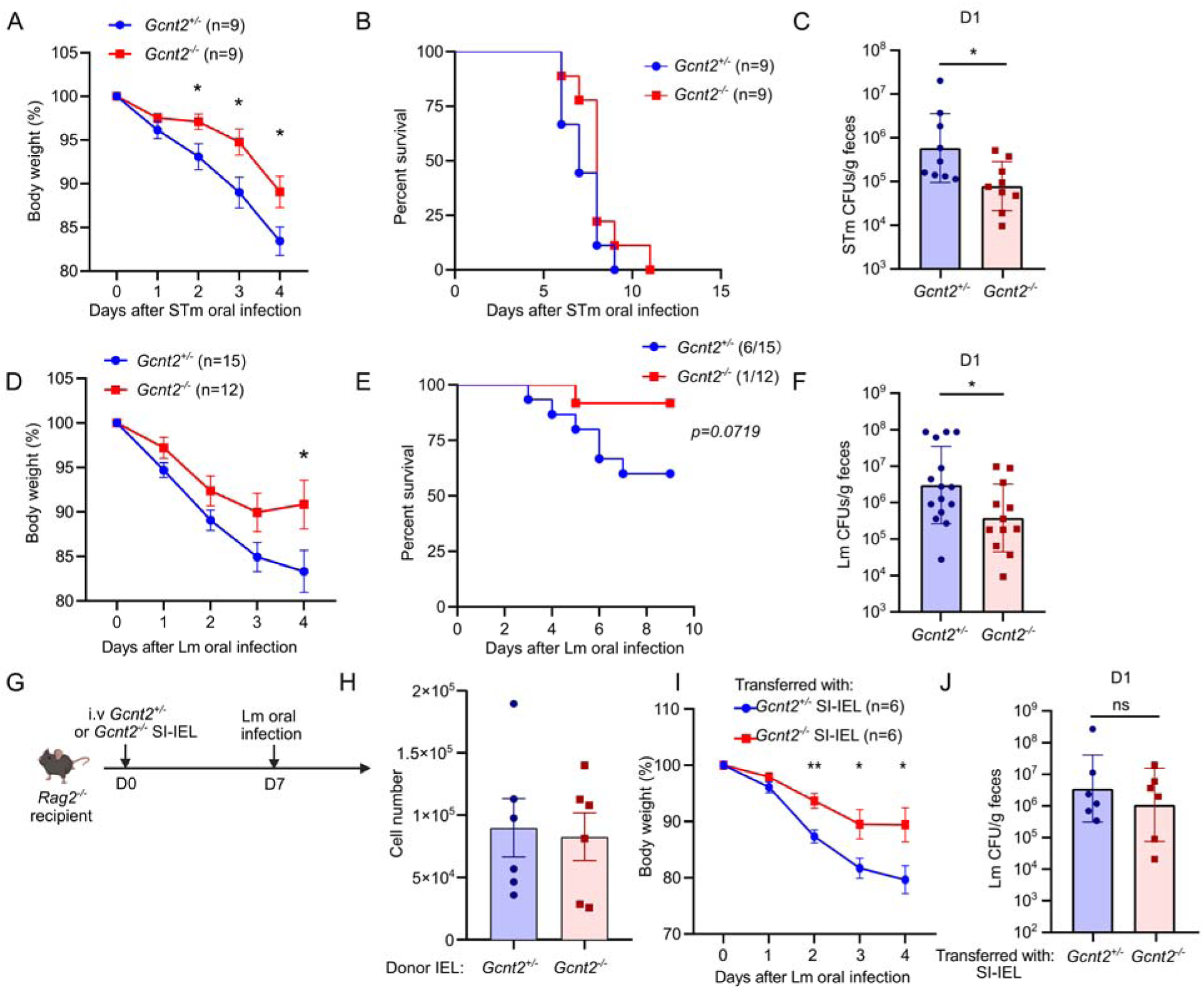
Loss of Gcnt2 confers resistance to oral bacterial infection. (A-C) Quantification of body weight loss (A), survival (B) and bacterial loads in feces at day 1 (C) after oral infection with Salmonella Thyphymurium (1×10^7^) in *Gcnt2^+/-^* and *Gcnt2^-/-^* mice. (D-F) Quantification of body weight loss (D), survival (E) and bacterial loads in feces at day 1 (F) after oral infection with *Listeria monocytogenes* (1×10^8^) in *Gcnt2^+/-^* and *Gcnt2^-/-^* mice. (G) SI-IE CD4^-^CD8αα^+^ T cells from *Gcnt2*^+/-^ and *Gcnt2*^-/-^ mice were transferred to *Rag2*^-/-^ recipients, followed by oral infection with *Listeria* at day 7 after cell transfer. (H) nIEL reconstitution detection at day 7 after cell transfer described in (G). (I-J) Body weight loss (I) and fecal *Listeria* loads at day 1 post infection (J) in recipient mice described in (G). Data in (A-C) were pooled from two independent experiments with n=9. Data in (D-F) were pooled from three independent experiments with n=12-15. Data in (H)-(J) are pooled from two independent experiments with n=6. Data in (A), (D), (H) and (I) are shown as mean ± SEM. Data in (C), (F) and (J) are show as geometric mean with geometric SD. Statistical analysis: unpaired t test (A, D and I); Mann-Whitney test (C, F and J); Log-rank (Mantel-Cox) test in (E). *p<0.05; **p<0.01; ns, not significant.

To directly test whether the absence of *Gcnt2* within IELs enhances host defense, we adoptively transferred enriched *Gcnt2*^+/-^ or *Gcnt2*^-/-^ CD4^-^CD8αα⁺ IELs into *Rag2*^-/-^ recipients and challenged them orally with *Lm* one week later (**Fig. 5G**). Reconstitution led to similar, low numbers of nIELs in the intestinal epithelium of the recipient mice (**Fig. 5H**). However, despite showing similar fecal bacterial burden, mice receiving *Gcnt2*^-/-^ IELs exhibited reduced weight loss, indicating that *Gcnt2* loss augments nIEL-dependent protection *in vivo* (**Fig. 5, I-J**).

We next asked whether GCNT2 similarly modulates induced CD8αβ⁺ IELs (Trm). Co-transfer of naïve *Gcnt2*^+/-^ and *Gcnt2*^-/-^ OT-I cells into WT recipients followed by oral *Lm*-*OVA* infection yielded Trm populations with comparable persistence and phenotype (Fig. S8, A-D). However, upon *ex vivo* SIINFEKL stimulation, *Gcnt2*^-/-^ Trm cells from the small intestine but not from the spleen, displayed enhanced degranulation (**Fig. S8E**), consistent with a tissue-specific impact of GCNT2 on T cells. *In vivo*, individual transfer of *Gcnt2*^+/-^ or *Gcnt2*^-/-^ OT-I cells followed by *Lm-OVA* immunization showed that *Gcnt2*-deficient Trm conferred superior protection against secondary *Yersinia*-*OVA* infection, reflected by lower fecal bacterial loads (**Fig. S8, F and G**). Together, these findings indicate that GCNT2 restrains effector activation in both natural and induced CD8⁺ IELs *in vivo*, thereby limiting their protective capacity during orogastric bacterial infection.

### Complete *Gcnt2* deficiency is associated with mild intestinal alterations and heightened responses to T cell–activating stimuli

We observed that Gcnt2^-/-^ mice exhibited reduced weight gain (**Fig. S9A**) and shortened colons (**Fig. S9B**), features potentially indicative of low-grade intestinal inflammation. In addition, we observed moderately increased expression of Lcn2 in both small intestine and colon and trends toward increased Il1b and Ifng transcripts (**Fig. S9C**). However, FITC-dextran assays revealed no increase in gut permeability (**Fig. S9D**), indicating intact barrier function. Furthermore, intestinal tissue organization and microbiota composition appeared unaltered in Gcnt2^-/-^ mice (**Fig. S9E-F and Fig. S10A-B**), excluding occurrence of overt inflammation.

Notably, *Gcnt2* deficiency on the OT-I background did not affect body weight or alter *Il1b* and *Lcn2* expression (**Fig. S9G and H**), suggesting that T cells with a diverse repertoire may contribute to the phenotypes observed in complete *Gcnt2*-deficient mice. Transfer of either CD8αα+TCRγδ+ IELs, which we deemed more likely to favor tissue damage, or thymic CD4-CD8-TCRγδ+ IEL precursors from *Gcnt2*^+/-^or *Gcnt2*^-/-^ mice into *Rag2*^-/-^ recipients did not reproduce the body weight differences observed between *Gcnt2*^+/-^ and *Gcnt2*^-/-^ mice (**Fig. S9I and J**). These results suggest either that CD8αα+TCRγδ+ IELs do not drive this phenotype, or that additional lymphoid populations and/or cellular interactions are required for its manifestation. Thus, while complete *Gcnt2* deficiency results in phenotypic alterations suggestive of mild gastrointestinal perturbation, the biological significance of these findings and the relative contribution of IELs remain uncertain.

We next asked whether GCNT2 can regulate responses to acute inflammatory stimuli. Anti-CD3 administration, which triggers rapid intestinal pathology via T cell activation (Merger et al., 2002; Radojevic et al., 1999; Xu et al., 2021), caused greater weight loss in *Gcnt2*^-/-^ mice (**Fig. S9K**). To model dietary-antigen–driven T cell activation, we delivered SIINFEKL orally to OT-I mice. As expected, OT-I but not WT mice showed transient weight loss and recovery (**Fig. S9L**). However, *Gcnt2*^-/-^ OT-I mice displayed increased weight loss (**Fig. S9M**) and higher *Lcn2* expression (**Fig. S9N**) at day 1, consistent with amplified intestinal inflammation following antigen encounter. Collectively, these findings indicate that GCNT2 influences basal physiological parameters as well as the consequences of stimulus-induced T cell activation, raising the possibility that loss of I-branched glycosylation may increase susceptibility to inflammation triggered by dietary, microbial, or other environmental cues.

### GCNT2 limits TCR signaling by glycosylating CD45

To identify the molecular targets through which GCNT2 modulates T cell activation, we probed total CD8α^+^ IEL lysates with biotinylated LEL. This revealed a prominent ∼200 kDa band in control but not *Gcnt2*^-/-^ IELs (**Fig. 6A**). Mass spectrometry identified CD45 as the only membrane-associated candidate among otherwise intracellular contaminants (**Fig. S11A**). CD45 immunoprecipitation followed by LEL blotting confirmed GCNT2-dependent glycosylation (**Fig. 6B**), consistent with earlier findings in B cells (Giovannone et al., 2018). Supporting this, overexpression of WT but not catalytically inactive *Gcnt2* (Zhang et al., 2011), enhanced CD45 glycan branching in *Gcnt2*^-/-^ OT-I splenocytes (**Fig. 6C**). Although LAMP2 was previously reported as a GCNT2 substrate in kidney tissue (Chen et al., 2005), neither LAMP1 nor LAMP2 showed GCNT2-dependent glycosylation in *Gcnt2* overexpressing cells (**Fig. S11B**).

**Figure 6.**
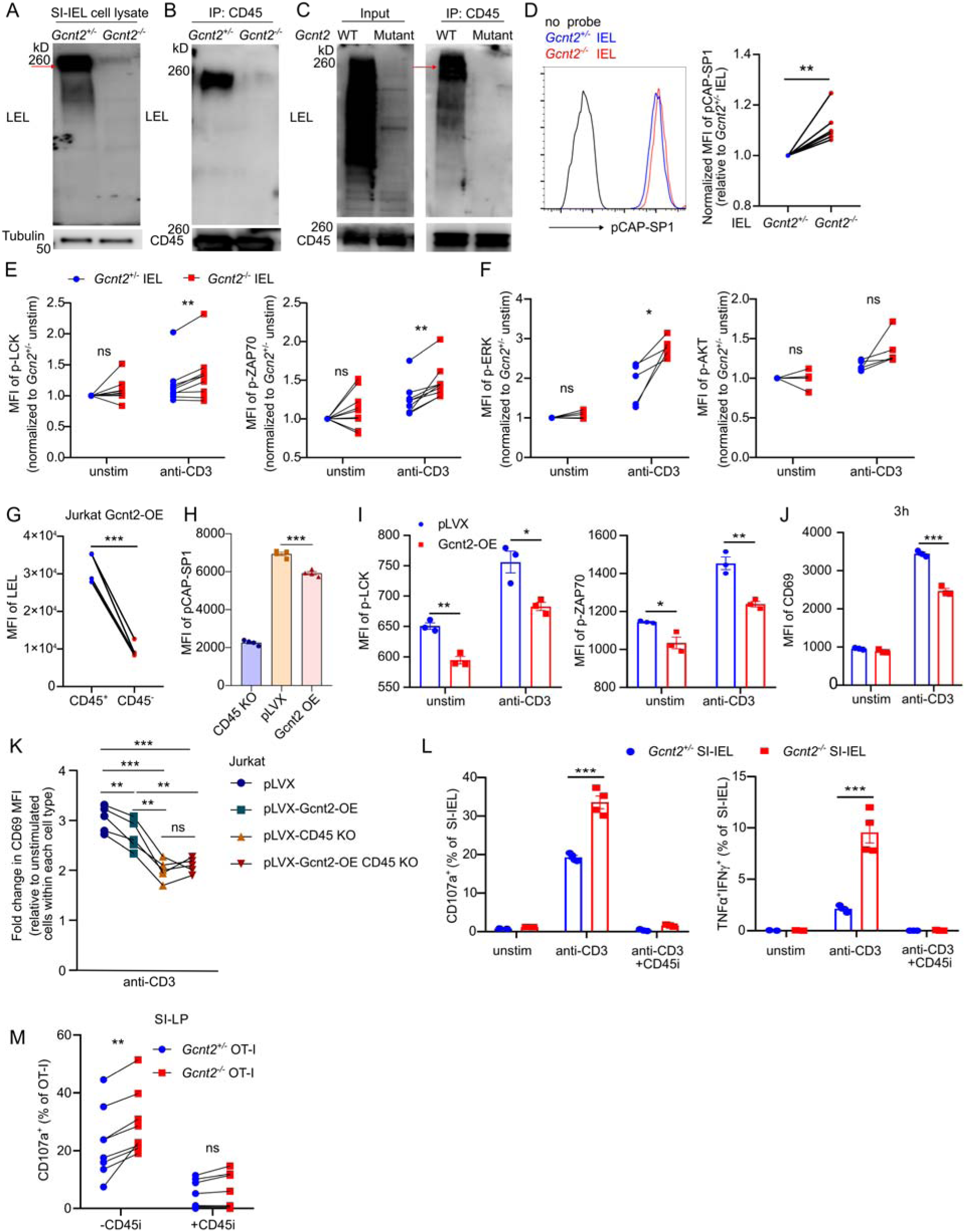
GCNT2 glycosylates CD45 to restrain TCR signaling. (A) Immunoblot using LEL to detect glycosylated proteins in total SI-IEL CD8α^+^ T cell lysates from *Gcnt2*^+/-^ and Gcnt2^-/-^ mice. The arrow indicates a potential dominant target. (B) CD45 was immunoprecipitated from *Gcnt2*^+/-^ and *Gcnt2*^-/-^ total SI-IEL CD8α^+^ T cell lysates, followed by LEL blotting to assess glycosylation. (C) Enriched naïve *Gcnt2*^-/-^ OT-I cells from spleen were *in vitro* activated and transduced with either WT or mutant *Gcnt2.* CD45 was immunoprecipitated from lysates, followed by LEL blotting to assess glycosylation. (D) FACS plot and normalized MFI of pCAP-SP1 in *Gcnt2*^+/-^ and *Gcnt2*^-/-^ SI-IEL CD4^-^CD8αα^+^ T cells. (E) Normalized MFI of phosphorylated LCK (Y394) and ZAP70 (Y319) at baseline or 5 min after plate coated anti-CD3 (5 µg/ml) stimulation in *Gcnt2*^+/-^ and *Gcnt2*^-/-^ SI-IEL CD4^-^CD8αα^+^ T cells. (F) Normalized MFI of phosphorylated ERK (T202/Y204) and AKT (S473) at baseline or 5 min after plate coated anti-CD3 (5 µg/ml) stimulation in *Gcnt2*^+/-^ and *Gcnt2*^-/-^ SI-IEL CD4^-^CD8αα^+^ T cells. (G) MFI of LEL staining in Gcnt2-overexpressing Jurkat cells with high or low CD45 expression generated by Crispr/Cas9. (H) CD45 phosphatase activity assay using pCAP-SP1 probe in CD45 KO, pLVX vector control and pLVX-*Gcnt2* overexpressing Jurkat cells. (I) MFI of phosphorylated LCK (Y394) and ZAP70 (Y319) at baseline or 10 min after anti-CD3 (5 µg/ml) stimulation in Jurkat cells expressing empty vector or *Gcnt2*. (J) MFI of CD69 expression at baseline or 3 h after anti-CD3 (5 µg/ml) stimulation in Jurkat cells expressing empty vector or *Gcnt2*. (K) Normalized MFI of CD69 expression in Jurkat cells under four conditions: pLVX control, pLVX-*Gcnt2* overexpression, CD45-deficient pLVX control, and CD45-deficient pLVX-*Gcnt2* overexpression. The cells were stimulated with anti-CD3 (5 µg/ml) for 3 h. (L) SI-IEL CD4^-^CD8αα⁺ T cells isolated from *Gcnt2*^+/-^ or *Gcnt2*^-/-^ mice were stimulated with plate-bound anti-CD3 (5 µg/ml) in the presence of CD107a antibody and Golgi inhibitor, with or without CD45 phosphatase inhibitor. After six hours, degranulation and cytokine production were analyzed by FACS. (M) *Gcnt2^+/-^* or *Gcnt2^-/-^* OT-I cells with different congenic markers were co-transferred into CD45.2 WT recipients at a 1:1 ratio, followed by oral infection with *Listeria-OVA*. At day 30 post infection, isolated cells from small intestine lamina propria were co-cultured with SIINFEKL-loaded CD45.2 splenocytes in presence or absence of CD45 phosphatase inhibitor. Data in (A), (B), (C), (G), (H), (I), (J) and (L) are one representative of two independent experiments. Data in (D) is pooled from six independent experiments. Data in (E) is pooled from nine independent experiments. Data in (F) and (K) are pooled from five independent experiments. Data in (M) is pooled from two independent experiments. Data are shown as mean ± SEM. Statistical analysis: paired t test for (D), (E), (F), (G), (K) and (M); unpaired t test for (H), (I), (J) and (L). *p<0.05; **p<0.01; ***p<0.001; ns, not significant.

CD45 serves as a key phosphatase controlling TCR signaling (Courtney et al., 2019). Using the CD45-specific fluorogenic substrate pCAP-SP1 (Szodoray et al., 2021) (**Fig. S11C**), we observed increased basal CD45 phosphatase activity in *Gcnt2*^-/-^ CD4^-^CD8αα⁺ IELs compared with controls (**Fig. 6D**). Consistently, *Gcnt2*^-/-^ CD4^-^CD8αα⁺ IELs exhibited increased phosphorylation of LCK at the activating residue Y394, as well as enhanced phosphorylation of ZAP70 following plate-bound anti-CD3 stimulation, indicating augmented proximal TCR signaling (**Fig. 6E**). In addition, phosphorylation of the downstream signaling molecule ERK, but not AKT, was also increased in *Gcnt2*^-/-^ CD4^-^CD8αα⁺ IELs (**Fig. 6F**).

To confirm the effect of CD45 glycosylation on TCR signaling, we turned to Jurkat T cells. Gcnt2 overexpression increased overall LEL staining, at least in part, in a CD45-dependent manner, as loss of CD45 significantly reduced the signal (**Fig. S11, D and E and Fig. 6G**). pCAP-SP1 dephosporylation was markedly compromised in CD45 KO Jurkat cells, confirming the specificity of this assay. Gcnt2 overexpression also reduced pCAP-SP1 signal, indicating that I-branching glycosylation directly impairs CD45 phosphatase activity (**Fig. 6H**). Reduced CD45 activity in Gcnt2-overexpressing cells correlated with lower basal and stimulated phosphorylation of LCK and ZAP70, proximal targets of TCR triggering, along with diminished CD69 induction (**Fig. 6, I and J**). As O-glycosylation is impaired in Jurkat cells due to a Cosmc (C1GALT1C1) deficiency, we verified our results using a second cells line, the murine T cell hybridoma B3Z. Similar to what observed in Jurkat, overexpression of Gcnt2 in B3Z cells led to increased glycosylation and decreased CD69 expression following anti-CD3 antibody (**Fig. S11F and G**).

CD45 has a dual and context-dependent role in T cells, as it can license LCK activation by dephosphorylating its inhibitory tyrosine residue, while also dampening TCR signaling by dephosphorylating phosphorylated signaling motifs within the TCR complex and other proximal signaling intermediates (Courtney et al., 2019). Consistent with the dual role of CD45 in TCR signaling, CD45 deficiency in Jurkat cells increased basal phosphorylation of LCK and ZAP70. However, following anti-CD3 stimulation, phosphorylation of LCK, ZAP70, and the downstream signaling molecule ERK was reduced in CD45 KO cells compared with control cells (**Fig. S11, H and I**). Thus, despite increased basal phosphorylation of proximal signaling molecules, CD45-deficient cells showed impaired signal propagation upon TCR stimulation.

We next asked whether the effect of Gcnt2 overexpression on Jurkat cell activation depended on CD45. In CD45-sufficient cells, Gcnt2 overexpression reduced anti-CD3-induced CD69 expression, consistent with **Fig. 6J**. In contrast, CD45 deficiency decreased CD69 induction and abolished the difference between control and Gcnt2-overexpressing cells (**Fig. 6K**). Together, these data indicate that GCNT2-dependent glycosylation restrains TCR signaling in model T-cell lines, at least in part through CD45.

We then examined the impact of CD45 inhibition in primary T cells. In CD8αα^+^ IELs, pharmacological inhibition of CD45 nearly abolished anti-CD3-induced activation, confirming the essential role of CD45 in TCR signaling but preventing us from assessing GCNT2-dependent differences in this setting (**Fig. 6L**). In intestinal OT-I Trm cells, CD45 inhibition attenuated, but did not completely abolish, SIINFEKL-induced activation. Under these conditions, the difference in CD69 induction between control and *Gcnt2*^-/-^ OT-I cells was lost (**Fig. 6M**). Together, these data indicate that GCNT2-mediated glycosylation dampens TCR signaling at least in part through modulation of CD45.

## Discussion

Intestinal intraepithelial lymphocytes (IELs) constitute a heterogeneous T cell network that integrates adaptive and innate features to preserve epithelial integrity. Positioned at the frontline of antigen exposure, IELs are primed for rapid cytotoxic responses yet must remain tightly restrained to avoid collateral tissue damage. Here, we identify GCNT2-dependent I-branching glycosylation as a checkpoint that attenuates TCR signaling, at least in part through CD45 modification, thereby maintaining IELs in a poised but self-controlled state.

At the molecular level, I-branching marks a subset of natural CD8αα⁺ IELs with reduced accessibility at stemness and TCR-signaling loci and increased accessibility at genes associated with tissue adaptation and immune regulation. These features parallel recent profiling that links reduced *Tcf7* and increased *Il2rb* or *Entpd1* to mature, self-restrained IEL states (Alonso et al., 2025; Yomogida et al., 2024). Because TCF1 marks precursor-like states, whereas CD122 and CD39 are hallmarks of tissue-adapted IELs (Alonso et al., 2025; Hioki et al., 2025; Moesta et al., 2020; Yakou et al., 2023), I-branching likely accompanies progression toward a more functionally mature, restrained phenotype. Whether mature I-Ag⁻ IELs can acquire I-branching or whether I-Ag⁺ cells represent a terminally adapted state remains unresolved.

Functionally, *Gcnt2* is not merely a marker of IEL identity. Its loss heightened IEL responsiveness, including effector cytokine production and degranulation, most prominently within the CD8αα^+^TCRγδ^+^ subset, and enhanced antibacterial protection *in vivo*. Although IELs have previously been implicated in defense against orogastric pathogens (Chawla et al., 2025; Chawla et al., 2024; Edelblum et al., 2015; Hoytema van Konijnenburg et al., 2017; Ismail et al., 2011), the mechanisms underlying this protection remain incompletely defined. However, the enhanced protective capacity associated with *Gcnt2* deficiency may come with trade-offs. *Gcnt2*^-/-^ mice exhibited reduced body weight gain, shortened colon length, and mildly increased expression of inflammatory markers. While these features are compatible with subclinical intestinal perturbation, we did not detect overt inflammation by histological or functional analyses. In addition, in models of direct T-cell stimulation, including anti-CD3 treatment and SIINFEKL gavage in OT-I mice, *Gcnt2*-deficient animals displayed increased body weight loss, which is commonly used as a readout of pathology in these settings.

These *in vivo* phenotypes should nevertheless be interpreted with caution. GCNT2-dependent glycosylation was detected not only in CD8αα^+^TCRγδ^+^ and CD8αα^+^TCRαβ^+^ IELs (Nie et al., 2022; Poussier et al., 2002), but also in other intestinal immune populations, including lamina propria Th17 cells and ILC3s. Thus, while our transfer experiments in *Rag2*^-/-^ mice support an IEL-intrinsic role for *Gcnt2* in protection from bacterial infection, the contribution of *Gcnt2* expression in distinct immune lineages to this as well as other in vivo phenotypes observed here remains to be elucidated.

Mechanistically, we identified CD45 as the predominant GCNT2-dependent glycosylation substrate in CD8αα^+^ IELs. Although GCNT2 is likely to modify additional, less abundant cellular proteins, our data show that loss of *Gcnt2* increases CD45 phosphatase activity at baseline and enhances TCR signaling following anti-CD3 stimulation in primary CD8αα^+^ IELs. Several factors may contribute to glycosylation-mediated restraint upon TCR triggering. Glycosylation modulates CD45 interaction with galectins, which can either enhance or inhibit CD45 activity (Earl and Baum, 2008; Szodoray et al., 2021) and even induce apoptosis in activated T cells (Earl and Baum, 2008; Nguyen et al., 2001). Because galectins preferentially bind highly branched poly-N-acetyllactosamine structures, a galectin lattice may sequester CD45 and limit its access to the TCR complex, thereby dampening synaptic signaling (Chen et al., 2007). Multiple galectins produced by activated T cells or intestinal epithelial cells can modulate TCR signaling through CD45 binding (Chabot et al., 2002; Chen et al., 2009; Earl et al., 2010; Okoye et al., 2020). In human GC B cells, GCNT2-dependent CD45 modification reduces Galectin-9 binding and relieves BCR inhibition (Giovannone et al., 2018) and because intestinal epithelial cells secrete Galectin-9 (de Kivit et al., 2017), similar regulation may occur in IELs.

The concept of branched N-glycans acting as a “glycan brake’’ on CD45 and TCR signaling may extend across species and diseases. In addition to enhanced antibacterial activity, our *in vitro* data suggest increased anti-tumor potential following *Gcnt2* loss. This is notable given the established contributions of γδ T cells to cancer immunity (Reis et al., 2022; Yakou et al., 2023) and the growing interest in glycan engineering, such as removal of MGAT5-dependent branches, to potentiate CAR T-cell responses (Azevedo et al., 2025; De Bousser et al., 2025). Human IELs differ phenotypically from murine IELs, and whether GCNT2-dependent glycosylation can also occur in human IELs is undetermined. Nevertheless, reduced N-glycan branching and impaired MGAT5 expression have been documented in IELs from ulcerative colitis patients (Dias et al., 2018; Dias et al., 2014; Pereira et al., 2020), and dysregulated IEL activity contributes to coeliac disease and IBD (Abadie et al., 2020; Ettersperger et al., 2016; Hue et al., 2004; Kornberg et al., 2023; Meresse et al., 2004; Meresse et al., 2006; Setty et al., 2015; Xu et al., 2025), underscoring the relevance of this regulatory axis in human disease. Within this framework, the observation that retinoic acid (RA) promotes *Gcnt2* expression and I-branching is of interest, as it links dietary vitamin A to IEL glycosylation. RA supports tolerance under steady-state conditions (Bakdash et al., 2015; Coombes et al., 2007; Kang et al., 2007; Mucida et al., 2007), consistent with the restraining function of I-branched glycans, although it can amplify inflammation when IL-15 is abundant, as in celiac disease (DePaolo et al., 2011). These insights raise the intriguing possibility that I-branching might be modulated through dietary or metabolic interventions.

IELs must operate under dual imperatives: rapid cytotoxicity against infected or transformed cells and steady-state protection of a fragile epithelium. Our findings support a model in which GCNT2-dependent I-branching contributes to this balance by limiting TCR-driven activation. Manipulating this pathway, nutritionally or therapeutically, may offer strategies to restrain intestinal inflammation while preserving barrier immunity. More broadly, our work highlights glycosylation as a tunable and underappreciated layer of immune regulation with relevance for infection, inflammation, and tissue homeostasis.

## Materials and methods

### Mice

All mice used in this study were on a C57BL/6J background. Wild-type (WT) C57BL/6J mice, CD45.1 WT mice, and CD45.2 OT-I mice were purchased from Charles River. *Rag2*^-/-^ mice were obtained from the laboratory of Dr. Stéphanie Hugues (University of Geneva). CD45.1 WT mice were crossed with CD45.2 OT-I mice to generate CD45.1 and CD45.1/2 OT-I mice, which were genotyped via Vβ5 FACS staining. Germ-free C57BL/6J mice were purchased from the University of Bern (Switzerland). *Gcnt2*^-/-^ mice were generated by the Transgenic Core Facility at the University of Geneva using CRISPR–Cas9–mediated genome editing. Two single-guide RNAs (sgRNAs) flanking the second and third exons of Gcnt2 were designed using the UCSC Genome Browser (https://genome.ucsc.edu/). The sgRNA target sequences were 5⍰-CCTGGTTCTTAGTGCAATGTCGT-3⍰ and 5⍰-ACCGTGAGGATCTGGCAACATGG-3⍰. crRNAs and tracrRNAs were synthesized by Integrated DNA Technologies (IDT) and resuspended in IDTE buffer. Equimolar amounts of crRNA and tracrRNA (200 pmol each) were annealed by heating to 95 °C for 5 min, followed by cooling at room temperature for 30 min. Each sgRNA was then complexed with recombinant Cas9 protein (1 µg/µl; IDT, #1081060) for 10 min at room temperature prior to microinjection into fertilized zygotes. sgRNA design, annealing, and zygote microinjection were performed by the Transgenic Core Facility at the University of Geneva.

Littermates of matched sex (both males and females) and age were used in all experiments. Mice were housed under specific pathogen–free conditions at the animal facility of the University of Geneva with a 12-h light/dark cycle. All animal procedures were approved and performed in accordance with the guidelines of the Geneva Cantonal Commission for Animal Experimentation (CCEA) and the Swiss Federal Food Safety and Veterinary Office (OSAV).

### Lymphocyte isolation

Spleen and thymus were mechanically dissociated through a 70-µm cell strainer to generate single-cell suspensions, followed by red blood cell lysis. OT-I cells were enriched using a CD8a⁺ T cell isolation kit (Miltenyi Biotec, 130-104-075).

For intestinal lymphocyte isolation, mesenteric fat and Peyer’s patches were carefully removed. Both the small and large intestines were opened longitudinally, cut transversely into ∼1-cm pieces, and transferred to 15 ml of IEL isolation buffer (HBSS without calcium and magnesium, supplemented with 20 mM EDTA [pH 8.0], 1 mM DTT, and 2% FBS). Tissues were incubated at 37 °C with shaking for 20 min. The supernatant was filtered through a 70-µm strainer to obtain a single-cell suspension enriched for IELs. This step was repeated twice.Remaining tissues were washed with PBS and digested in lamina propria digestion buffer (RPMI 1640 containing 2 mg/ml collagenase IV [Worthington], 10% FBS, 10 µg/ml DNase I, and 5 mM CaCl₂) at 37 °C with shaking for 40 min. The supernatant was collected and filtered through a 100-µm strainer to generate a single-cell suspension. Lymphocytes from both the epithelial and lamina propria fractions were further enriched using 40% Percoll (GE Healthcare).

### Flow cytometry and *in vitro* stimulation

Cells were incubated with anti-mouse CD16/32 antibody for 20 min on ice, followed by surface staining with the indicated antibodies for 30 min on ice in FACS buffer (PBS containing 5 mM EDTA and 1% FBS). The following antibodies (BioLegend) were used for surface staining: CD3 (17A2), CD8a (53-6.7), CD4 (GK1.5), CD8b (YTS156.7.7), CD19 (6D5), CD45 (30-F11), CD45.1 (A20), CD45.2 (104), TCRγδ (UC7-13D5), TCRβ (H57-597), CD5 (53-7.3), H-2K⍰ (AF6-88.5), CD44 (IM7), PD-1 (29F.1A12), CD122 (TM-β1), CD107a (1D4B), CD39 (Duha59), TCR Vγ1.1+1.2 (4B2.9), TCR Vγ2 (UC3-10A6), TCR Vγ3 (536), TCR Vγ4 (49.2), TCR Vγ7 (F2.67), LAG3 (C9B7W), human CD69 (FN50) and human CD45 (HI30).

OSK-14 antibody (a gift from the Japanese Red Cross, Osaka) was used at a 1:10 dilution, followed by staining with anti-human IgM secondary antibody (MHM-88). Biotinylated LEL and PHA-L lectin (Vector Laboratories) was used at a 1:1500 dilution to minimize binding-induced cell death, followed by staining with fluorophore-conjugated streptavidin (Invitrogen). For Galectin-9 binding, IELs were firstly incubated with recombinant human Galectin-9 (Biolegend, 1 µg/ml), followed by staining with PE anti-human/mouse Galectin-9 antibody (W21101F, Biolegend).

For intracellular cytokine staining, cells were stimulated with either SIINFEKL peptide or plate-bound anti-CD3 antibody (BioLegend) in the presence of protein transport inhibitors (eBioscience) or CD107a antibody for 4 h at 37 °C, unless otherwise indicated. Cells were then stained for surface markers and fixable viability dye (BioLegend), fixed and permeabilized using the True-Nuclear Transcription Factor Buffer Set (BioLegend), and stained with antibodies against cytokines.

For phosphorylation analysis, cells were stimulated for the indicated time points and fixed in 2% paraformaldehyde in PBS for 30 min at 37 °C. Cells were then pelleted, chilled on ice for 1 min, and permeabilized with 90% methanol for 30 min on ice. After washing with ice-cold incubation buffer (PBS containing 0.5% BSA), cells were blocked for 10 min at room temperature and stained with antibodies diluted in incubation buffer for 1 h at room temperature. Antibodies used included RORγt (AFKJS-9), FOXP3 (MF-14), granzyme B (QA16A02), granzyme A (3G8.5), IFN-γ (XMG1.2), TNF-α (MP6-XT22), phospho-LCK (Tyr394, A18002D), phospho-ZAP70/Syk (Tyr319, Tyr352, Zap70Y319-A3), phospho-ERK1/2 (Thr202/Tyr204, 4B11B69) and phospho-AKT (Ser473, A21001C).

Flow cytometry data were acquired on an LSRFortessa (BD Biosciences) and analyzed using FlowJo software. Cell sorting was performed using a BD Aria cell sorter.

For antigen-specific restimulation of Trm cells, CD45.2⁺ splenocytes from WT mice were pulsed with varying concentrations of SIINFEKL peptide for 30 min at 37 °C. Trm-containing IEL or lamina propria cells were then co-cultured with peptide-loaded splenocytes in the presence of CD107a antibody and protein transport inhibitors for 4 h at 37 °C.

### *Listeria-OVA* infection model

*Listeria monocytogenes OVA*-expression strain was constructed on the *10403S* background (Pope et al., 2001). *Lm*-*OVA* was used for Trm generation in this study. Bacteria were cultured overnight in brain heart infusion (BHI) broth at 37℃, then sub-inoculated and grown until early log phase (OD = 0.2-0.4) for enumeration. 16 hours before infection, mice were gavaged with Streptomycin (20 mg/mouse) as previously reported (Becattini et al., 2020; Cheng and Becattini, 2024) and feed was withdrawn from the cages. Simultaneously, freshly isolated 1.5×10^5^ CD45.1 WT OTI cells or *Gcnt2*^+/-^ and *Gcnt2*^-/-^ OTI cells with different congenic markers (1:1 mix) were co-transferred to recipient mice.

To administer *Lm*-OVA orally, we took advantage of an approach described elsewhere which avoids accidental systemic delivery of the bacterium via mechanical trauma caused by the gavage needle(Bou Ghanem et al., 2013; Louie et al., 2019). On day 0 (following 16h of fasting), mice were fed with bread cubes containing 5×10^7^ CFUs of *Lm*-OVA and melted butter. Following bread consumption, feed was placed back into the cages.

### *Salmonella, Listeria, Yersinia-OVA* infection model

*Salmonella Typhimurium* (SL1344) was a gift from Dr. Wolf-Dietrich Hardt, ETH, Zurich. For experiments, single colonies were picked from LB plates into LB medium and cultured with shaking for around 4 hours. Mice were gavaged with 1×10^7^ CFUs of STm in PBS. Feces were collected, weighted, resuspended in PBS and plated on LB agar plates with Chloramphenicol (20 µg/ml) at different days after infection. WT *Listeria monocytogenes* was cultured in BHI medium, and mice were gavaged with 1×10^8^ CFUs. Feces were collected and plated on BHI agar plates with Streptomycin (100 µg/ml) and Nalidixic Acid (50 µg/ml).

Yersinia Pseudotuberculosis OVA-expressing strain (Yptb-OVA, MB323) is a gift from Dr. Peter Dube, Department of Microbiology & Immunology, The University of Texas Health Science Center San Antonio, San Antonio, TX, United states of America. The strain produces a fusion protein containing the natural Yersinia antigen YopE (residues 1-138) fused with chicken ovalbumin (residues 247-355) and was previously shown to elicit OVA-specific CD8 T cell responses(Gonzalez-Juarbe et al., 2017). Single colony was inoculated in LB medium overnight at room temperature. Mice were gavaged with 3×10^8^ CFUs of MB323. Feces were collected and plated on LB agar plates with Yersinia Selective Supplement (Sigma).

### Bulk RNA-sequencing

To exclude a potential influence of circulating T cells, 3-5 min prior to sacrifice, mice were intravenously injected with anti-CD45 antibody. Resident cells or previously transferred OTI cells were sorted from the indicated organs, followed by RNA extraction (Microprep RNeasy, Qiagen). Library preparation was performed using Smarter Nextera reagents, sequencing was performed with Illumina NovaSeq (Genomics Platform of iGE3, University of Geneva). FastQ reads were mapped to the ENSEMBL reference genome (GRCm39.103) using STAR version 2.4.0j(Dobin et al., 2013) with standard settings, except that any reads mapping to more than one location in the genome (ambiguous reads) were discarded (m = 1). A unique gene model was used to quantify reads per gene. Briefly, the model considers all annotated exons of all annotated protein coding isoforms of a gene to create a unique gene where the genomic region of all exons are considered coming from the same RNA molecule and merged together. All reads overlapping the exons of each unique gene model were reported using featureCounts version 1.4.6- p1(Liao et al., 2014). Gene expressions were reported as raw counts and in parallel normalized in RPKM in order to filter out genes with low expression value (1 RPKM) before calling for differentially expressed genes. Library size normalizations and differential gene expression calculations were performed using the package edgeR(Robinson et al., 2010) designed for the R software. Only genes having a significant fold-change (Benjamini-Hochberg corrected p-value < 0.05) were considered for the rest of the RNAseq analysis. For reanalysis of published RNA-seq dataset, DESeq2 was used to do differential gene expression analysis(Love et al., 2014).

### RT-qPCR

RNA from purified cells was extracted according to the manufacturer’s protocol (Microprep RNeasy, Qiagen). A piece of small intestine or colon were homogenized in Trizol (Invitrogen), and total RNA was extracted using chloroform separation and isopropanol precipitation, followed by ethanol wash and DNA removal using DNA-free kit (Invitrogen). Subsequently, cDNA was synthesized by using QuantiTect Reverse Transcription Kit (Qiagen). RT-qPCR was performed using PowerUp SYBR green (Thermo Fisher Scientific). The following primers were used: m*Gcnt2* isoform A-F: TACGTGATCAGAACCACGGCTCT; m*Gcnt2* isoform A-R: CCATCCACTTCACAGCTCTGAG; m*Gcnt2* isoform B-F: AGTCCTGATGAGCACTTCTGGGTGA; m*Gcnt2* isoform B-R: AGTTCAAATTTGTTAGCAAACAGG; m*Gcnt2* isoform C-F: TGAATGACGAAAGGGCCATTG; m*Gcnt2* isoform C-R: CAGCCACTGCAAGTCTCCG; m*Actin*-F: TACCACCATGTACCCAGGCA; *mActin*-R: CTCAGGAGGAGCAATGATCTTGAT; *mGzma*-F: TGCTGCCCACTGTAACGTG; *mGzma*-R: GGTAGGTGAAGGATAGCCACAT; *mGzmb*-F: CCACTCTCGACCCTACATGG; *mGzmb*-R: GGCCCCCAAAGTGACATTTATT; *mTcf7*-F: AGCTTTCTCCACTCTACGAACA; *mTcf7*-R: AATCCAGAGAGATCGGGGGTC; *mIfng*-F: CAGCAACAGCAAGGCGAAAAAGG; *mIfng*-R: TTTCCGCTTCCTGAGGCTGGAT; *mTnfa*-F: CTACCTTGTTGCCTCCTCTTT; *mTnfa*-R: GAGCAGAGGTTCAGTGATGTAG; *mLcn2*-F: TGGCCCTGAGTGTCATGTG; *mLcn2*-R: CTCTTGTAGCTCATAGATGGTGC; *mCcl2*-F: GCTACAAGAGGATCACCAGCAG; *mCcl2*-R: GTCTGGACCCATTCCTTCTTGG; *mIl1b*-F: ACGGACCCCAAAAGATGAAG; *mIl1b*-R: TTCTCCACAGCCACAATGAG.

### FITC-Dextran permeability assay

Mice were deprived of food for 4 hours. FITC-Dextran (MW3000-5000, Sigma) was dissolved in PBS at a concentration of 100 mg/ml and orally gavaged to mice at a dosage of 10 mg/mouse. Food was replaced immediately following the gavage. After 4 hours, serum was be collected, and the concentration of FITC in serum was determined using SpectraMax Paradigm Multi-Mode Microplate Reader (Molecular Devices).

### IEL precursor cell transfer

Thymus from 4-week old mice were meshed through a 70 µm strainer to obtain single-cell suspensions. CD4^+^ and CD8α^+^ positive selection kits (Miltenyi) were used to remove CD4^+^CD8^-^, CD4^+^CD8^+^ and CD4^-^CD8^+^ populations. The double negative cells containing IEL precursors were enriched in the flow-through. Both CD4^-^CD8^-^TCRβ^+^ and CD4^-^CD8^-^TCRγδ^+^ cells were sorted and intravenously transferred into *Rag2*^-/-^ recipient mice for experiments evaluating I-Ag expression or retinoic acid receptor antagonist treatment. Only CD4^-^CD8^-^TCRγδ^+^ cells were sorted and intravenously transferred into *Rag2*^-/-^ recipient mice for the experiments shown in Fig S9 to determine whether *Gcnt2*^-/-^ CD8αα^+^TCRγδ^+^ IELs contribute to reduced body weight gain in mice.

### Retinoic acid and retinoic acid receptor antagonist treatment *in vivo*

Retinoic acid (R2625, Sigma) was dissolved in DMSO (stock concentration, 50 mg/ml), further diluted in corn oil before oral gavage (200 µg/mouse/day). RA was supplemented every other day for three weeks. Mice were injected intraperitoneally with 1mg/kg pan-RAR antagonist AGN194310 (Sigma) every other day. Vehicle (DMSO+PBS) was used for control mice injection.

### Antibiotics treatment *in vivo*

For the oral antibiotic treatment experiment, mice were provided drinking water supplemented with a cocktail of broad-spectrum antibiotics for one month. The antibiotics used included ampicillin (0.5 g/L, Applichem), metronidazole (0.5 g/L, MP Biomedicals), neomycin (0.5 g/L, ThermoFisher) and vancomycin (0.5 g/L, ThermoFisher). Feces were collected and plated on Columbia Blood Agar Plates (BD) in an anaerobic chamber to assess the efficiency of microbiota depletion.

### Retroviral transduction of OT-I cells

Plat-E cells were seeded in 12-well plates at a density of 1.5 × 10⁵ cells per well 12 h prior to transfection. Cells were transfected with pCL-Eco (0.5 µg) together with pMSCV-GCNT2-GFP, pMSCV-GCNT2-mutant-GFP, pMSCV-GFP, or pMSCV-dnRARα-GFP (1 µg; constructs provided by the laboratory of Dr. Laura Mackay, University of Melbourne) using TransIT-LT1 Transfection Reagent (Mirus). Retroviral supernatants were collected 48 h post-transfection, filtered through a 0.45-µm filter, and concentrated by centrifugation at 14,000 rpm for 2 h.

Primary T cells were cultured in RPMI 1640 (Gibco) supplemented with 10% heat-inactivated FBS (Gibco), 2 mM L-glutamine (Gibco), 1 mM sodium pyruvate (Gibco), non-essential amino acids (Gibco), 100 U/ml penicillin, 100 µg/ml streptomycin, and 50 µM 2-mercaptoethanol (Gibco). Purified OT-I cells were stimulated with plate-bound anti-CD3 and anti-CD28 antibodies (5 µg/ml each; BioLegend) for 24 h, then spin-transduced with retroviral supernatant in the presence of polybrene (10 µg/ml; Invitrogen) at 1,800 rpm and 32 °C for 2 h.

Six hours after transduction, cells were transferred to fresh plates and expanded for an additional 2 d in the presence of IL-2 (10 ng/ml; BioLegend) prior to adoptive transfer into WT recipient mice, which were subsequently infected orally with *Listeria monocytogenes*–OVA (Lm-OVA).

### *In vitro* activation and stimulation of OT-I cells by retinoic acid

To assess short-term effects of retinoic acid (RA) or cytokines on *Gcnt2* induction, purified OT-I cells were stimulated with plate-bound anti-CD3 and anti-CD28 antibodies (5 µg/mL each) for 3 days, collected, resuspended in fresh medium, and restimulated with the indicated stimuli for 6 hours. To evaluate long-term effects, purified OT-I cells were stimulated with plate-bound anti-CD3 and anti-CD28 in the presence of IL-2 (10 ng/ml) and the indicated stimuli for 3 days, then transferred to fresh plates with newly supplied IL-2 and stimuli. On day 6, cells were harvested and analyzed for *Gcnt2* mRNA expression by RT-qPCR. Cells were treated with IL-15 (10 ng/ml), RA (10nM) or TGF-β (10 ng/ml) as indicated.

### Immunofluorescent staining and quantifications

Tissues were harvested, fixed in 4% PFA overnight, embedded into OCT. Cryosections were thawed for 20 minutes at room temperature, fixed in PFA 4% for 5 minutes. All samples were washed with PBS and permeabilized with 0.3% Triton-X-100 in PBS, followed by 30 minutes blocking (5% donkey serum, 0.5% BSA, and 0.3% TritonX-100 in PBS), overnight incubation with primary antibody (CD8α, #14-0195-82, eBioscience, 1:200; E-cadherin, #AF478, R&D Systems, 1:300) and secondary antibody staining for 1 hour. Slides were mounted in Fluoromount-G mounting medium with DAPI (Invitrogen, #00-4959-52). A NanoZoomer S60 slide scanner (Hamamatsu) with a 20x objective was used to scan the slides. Images were analyzed with NDP.view2 (Hamamatsu), and open-source software Qupath(Bankhead et al., 2017). CD8α^+^ cells were isolated using object classifier and their abundance was normalized to total villi area (at least 10 villi per samples were analyzed). The distance of CD8α^+^ from villi base was manually annotated in Qupath.

### ATAC-seq

For ATAC-seq, analyses were performed as recently reported (Schwaemmle et al., 2025). 60,000 cells were pelleted at 500 rcf, at 4⍰°C for 5⍰min. Supernatant was removed, cells were resuspended in 50⍰µl RSB buffer (10 mM Tris-HCl pH7.4, 10 mM NaCl, 3 mM MgCl_2_) containing 0.1% Tween-20 and 0.1% NP40 and incubated on ice for 3⍰min. Lysis was washed out with 500⍰µl RSB buffer containing 0.1% Tween-20. Samples were centrifuged for 10⍰min at 4⍰°C and 500 rcf. Supernatant was removed and pellets resuspended in 50⍰µl transposition mix (25⍰µl TD buffer (Diagenode, C01019043-1000), 2.5⍰µl Tagmentase (Diagenode, C01070012-200), 22.5⍰µl H_2_O). Samples were incubated for 30⍰min at 37⍰°C at 1000⍰rpm. DNA was cleaned up using the MinElute kit (Qiagen, 28004). Eluted DNA was amplified for 10 cycles, adding Nextera Ad PCR primers. DNA was cleaned up using the MinElute kit (Qiagen, 28004). Library molarity and quality were assessed by Qubit and Tapestation (DNA High sensitivity chip). Libraries were sequenced on a NovaSeq 6000 Illumina sequencer with paired-end 50 settings. Sequencing reads were trimmed using cutadapt 3.5 (adapters -m 20 -O 5 -p). Reads were aligned to the mm10 genome using bowtie2 v2.4.4 (--maxins 2000 -p 8 -N 1). Alignments were filtered using samtools v1.12 (view -h -F 1796 -q 20). ATAC peaks were called using MACS2 v2.7.1 (callpeak --format BAMPE --call-summits --shift −75 --extsize 150 --keep-dup all -B --SPMR -g mm -t ATAC.bam). Peaks from all replicates within 1⍰kb were merged using bedtools v2.30, filtered against the mouse blacklist, and the numbers of reads from each replicate overlapping with the peak set was counted using featureCounts v2.0.3 (-T 4 -O -p -a -t exon -g gene_id). Differential ATAC peak calls were determined using the R software DESeq2 after pre-filtering peaks with low counts (rowSums⍰>⍰100), with significance cut-offs set at FDR⍰<⍰0.05 and fold change > 1.5 in either direction. Output data were plotted with ggplot2. For IGV genome browser snapshots, average of replicates genome coverage files (bigwig) was generated with deepTools v3.5.2 (bamCoverage --binSize 10 --normalizeUsing RPKM --ignoreForNormalization chrM and bigwigAverage for merging replicates). Genomic features of ATAC peaks were determined using the R software ChIPSeeker v1.32.1. deepTools v3.5.2 was used to produce heat map of published SMARCA4 ChIP-seq mean read density across ATAC. GREAT software was used to annotate ATAC peaks with genes to determine gene ontology terms associated with changes in accessibility.

### 16S rRNA sequencing

The protocol was previously reported elsewhere (Czauderna et al., 2025). DNA from fecal samples was extracted using PureLink Microbiome DNA Purification Kit (Invitrogen) according to manufacturer’s protocol. V4 amplicons of the 16S rRNA gene were generated using universal bacterial primers containing barcode and Illumina adapters according to our established protocols (Czauderna et al., 2025). Duplicates of each amplicon were mixed, quantified with Qubit (Invitrogen), and pooled at equivalent concentrations. The final library was purified using GeneJET PCR purification kit (Thermo Scientific) and sequenced on the MiSeq Illumina platform (Illumina MiSeq V2 500 cycles, 250 pair-end) at Lausanne Genomic Technologies Facility. Paired-end raw reads were demultiplexed and processed using the QIIME2 pipeline (Bolyen et al., 2019). Briefly, sequences were trimmed with cutadapt algorithm, paired ends were merged with vsearch algorithm, quality-filtered using q-score method, and chimera-checked. Greengenes2 2022.10 database was used to construct the reference sequence file, on which the Naïve Bayes classifier was trained and then used to perform taxonomic assignment to the species level using representative sequences from each OTU.

### *In vitro* cytotoxicity assay

Sorted CD4^-^CD8αα^+^ *Gcnt2^+/-^* or *Gcnt2^-/-^* IELs were co-cultured with 1×10^4^ CFSE-labelled YAC1 or A20 cells at varying effector to target ratios in 96-well flat-bottom plate pre-coated with anti-CD3 (5 µg/ml). 24 hours later, cells were harvested, stained with a viability dye and target cell death was assessed by FACS.

### CD8 T cell depletion *in vivo*

Mice were injected intraperitoneally with anti-mouse CD8α antibody (100 µg per dose, BioXcell, #BE0061) every other day for a total of three doses prior to *Listeria* oral infection. Peripheral blood was collected the day before infection to assess CD8α⁺ T cell depletion efficiency. At day 5 post-infection, spleen and small intestine were harvested to confirm CD8α⁺ T cell depletion.

### IEL transfer

1×10^6^ sorted CD4^+^CD8αα^+^ *Gcnt2^+/-^* or *Gcnt2^-/-^* IELs were intravenously transferred to *Rag2^-/-^* recipient mice. One week after reconstitution, mice were orally gavaged with 1×10^8^ *Listeria* CFUs.

### Anti-CD3 antibody intraperitoneal injection model

Mice were injected i.p. with anti-CD3 antibody (BE0001-1, 25 μg, BioXCell) in 200 µl PBS, and the body weight and morbidity were monitored.

### SIINFEKL peptide gavage model

OT-I mice were orally gavaged with 100 µg SIINFEKL peptide (GenScript) in 200 µl PBS, and body weight was monitored thereafter.

### Western Blot and immunoprecipitation

Cells were lysed in Pierce IP lysis buffer (#87787, ThermoFisher) for 30 min on ice. Lysates were centrifuged at 4°C, 13.000 rcf for 10⍰min and supernatant transferred to new tubes. Samples were mixed with 4x NuPAGE LDS Sample Buffer (#NP0007, ThermoFisher) and boiled for 10 min at 70⍰°C. Then, samples were loaded onto mPAGE 4-12% Bis-Tris gel (MP41G12, Millipore) and run for 1 hour at 100 V in running buffer (#NP0001, Life technologies). Transfer was conducted using Trans-Blot Turbo Transfer System according to the manufacturer’s instructions (Biorad). Membranes were firstly blocked in 5% BSA in TBS-T (0.1% Tween20, #1706435, Biorad), and then incubated with primary antibody (biotinylated LEL, #B1175-1, VectorLab, 1:100; CD45, #14-0451-82, Invitrogen, 1:2000; CD107a, #sc-20011, Santa Cruz, 1:2000; CD107b, #sc-19991, Santa Cruz, 1:2000) overnight at 4⍰°C. Then, after wash, membranes were incubated with secondary antibody (Streptavidin-HRP, anti-mouse HRP or anti-rat HRP) for 1 hour at room temperature. Membranes were imaged on an Odyssey DLx imager (LICORbio). For immunoprecipitation, 30 µL of cell lysate was reserved as input, and the remaining lysate was incubated with CD45, CD107a or CD107b antibody (1 µg/sample) at 4 °C overnight with rotation. Protein A/G magnetic beads (25 µL; #88802, Thermo Fisher) were washed three times with Pierce IP lysis buffer, mixed with the antibody–lysate complex, and incubated for 2 h at 4 °C with rotation. Beads were then washed three times with lysis buffer, resuspended in loading buffer, and boiled to elute bound proteins. The beads were separated using a magnetic stand (#12321D, Invitrogen), and supernatants were collected for SDS-PAGE and immunoblotting as described above.

### Mass spectrometry

WT IEL cell lysate was loaded onto the gel, followed by Coomassie staining and wash by solution containing ethanol and acetic acid. The band with expected size was cut and sent for mass spectrometry in the platform of Functional Genomics Center Zurich. Gel bands were cut in small pieces and washed with 100 mM NH4HCO3/50% acetonitrile (2x) and acetonitrile (1x). All supernatants were discarded. For the digestion, the gel pieces were covered with a buffered trypsin solution at pH 8 (10 mM Tris/2 mM CaCl2). Samples were enzymatically digested. Supernatants were collected and the remaining peptides were extracted from the gel bands with 0.1% TFA / 50% acetonitrile. Both supernatants were combined and dried. The sample was further digested by PNGase F (New England lab) overnight at 37°C. The digested sample was dried by SpeedVac. The digested samples were dissolved in aqueous 3% Acetonitrile with 0.1% formic acid, and the peptide concentration was estimated with the Lunatic UV/Vis absorbance spectrometer (Unchained Lab). Peptides were separated on a M-class UPLC and analyzed on a Q-Exactive mass spectrometer (Thermo). The acquired MS data were processed using the Maxquant search engine(Cox and Mann, 2008).

### *Gcnt2* overexpression in Jurkat cells

293T cells were seeded in 12-well plates at 1.5 × 10⁵ cells per well 12 hours prior to transfection with pSPAx2 (0.5 µg), pMD2.G (0.5 µg), and either pLVX-GFP or pLVX-GCNT2-GFP (1 µg) using TransIT-LT1 transfection reagent (Mirus). Lentiviral supernatants were collected 48 hours post-transfection, filtered through a 0.45 µm membrane, and concentrated by centrifugation at 14,000 rpm for 2 hours. Jurkat cells were then spin-transduced with the concentrated lentivirus, and GFP⁺ cells were purified by flow cytometric sorting.

### Crispr/Cas9 gene editing of CD45 in Jurkat cells

sgRNA targeting human Cd45 (5’-GTATTTGTGGCTTAAACTCTTGG-3’) was synthesized from IDT. 2 µl (2 µM) crRNA was mixed with 2 µl (2 µM) tracRNA and 6 µl Duplex buffer, followed by 95°C 5 min and 10 min at room temperature. 1.5 µl annealed sgRNA was mixed with 1.5 µl Cas9 protein at room temperature for 20 min. 5×10^5^ Jurkat cells were resuspended in buffer R (#MPK10025, Invitrogen), and electroporated using the Neon Transfection System (Invitrogen), with the program: 1600 V, 10 ms, 3 pulses.

### CD45 phosphatase activity assay

A cell-permeable fluorogenic phosphorylated coumaryl amino propionic acid peptide (pCAP-SP1) specific for CD45 phosphatase activity was designed taking advantage of peptide sequences of known CD45 substrates, and previously validated (Stanford et al., 2012). This peptide contains phosphotyrosine. Once entering into cells, the peptide can be dephosphorylated by protein tyrosine phosphatases, and the resulting cell fluorescence (Pacific Blue) can be monitored by FACS. Cells were harvested and washed in RPMI medium (without phenol red) supplemented with 0.5% FBS and equilibrated to 37°C before incubation with pCAP-SP1 (final concentration, 10 µM) for 15 min at 37°C. Cells were washed in FACS wash buffer (Ca^2+^/Mg^2+^ free PBS, supplemented with 2% FBS, 10 mM sodium orthovanadate and 30% H_2_O_2_) to inhibit tyrosine phosphatase activity. Cells were then further washed and analyzed by FACS.

### Statistical analysis

Data are plotted showing the mean ± SEM. Statistical analysis was performed in Prism 8 with: unpaired t test, paired t test, Mann-Whitney test or Log-rank (Mantel-Cox) test as indicated. *p<0.05; **p<0.01; ***p<0.001; ****p<0.0001; ns, not significant.

## Data availability

Bulk RNA-seq and ATAC-seq data generated in this study have been deposited on Zenodo (DOI : 10.5281/zenodo.17391550), 16s rRNA sequencing data has been deposited on Zenodo (DOI: 10.5281/zenodo.19581417) and they are publicly available as of the date of publication. Sequencing data were analyzed using standard pipelines. This paper does not report new original code. All original data presented in this study are available and can be provided upon request.

## Acknowledgements

We thank members of the Becattini lab, Dr. Christoph Scheiermann and Dr. Mikael Pittet for fruitful discussion. We thank Dr. Petra Bacher, Dr. Georg Gasteiger and Dr. Sigrun Stulz for critical insight and experimental approaches not included in the final manuscript. We thank Dr. Stéphanie Hugues for providing *Rag2^-/-^*mice. We thank Dr. Fabrizio Thorel at the Transgenic core facility of the University of Geneva for support in generating *Gcnt2* KO mice. We thank the FACS facility and animal facility at University of Geneva for their assistance. We thank Dr. Peter Dube for providing the MB323 *Yersinia pseudotuberculosis* strain. We thank Dr. Laurance Genton Graf and Dr. Yves Dupertuis for sharing Germ-free mice. We thank Dr. Camila Jandus and Ziyang Su for sharing YAC1 cell line, and Dr. Federico Simonetta and Sisi Wang for sharing A20 cell line. This work was supported by an Eccellenza Professorial Fellowship (PCEFP3_187018) by the Swiss National Science Foundation and a Project Funding Young Researcher from the Helmut Horten Stiftung to S.B. S.B. is further supported by a Starting Grant (TMSGI3_211235) by the Swiss National Science Foundation. M.E. is supported by an ARC DECRA Fellowship (503658). L.K.M. is supported by an NHMRC Leadership Investigator Fellowship (GNT2034209). N.C. is supported by an ERC Advanced Grant ‘GlycoCAR’ as well as by a research grant from the Fund for Scientific Research-Flanders (FWO, G050420N). Schematic depitcions throughout this manuscript were created with BioRender.

## Author contributions

L.C. and S.B. conceived and designed the study. L.C. conducted the experiments, with assistance from N.B. and A.C. L.B. performed CD8 tissue staining under the supervision of T.P. H.S. and S.Braun performed sample preparation and data analysis for ATAC-seq. S.L. analyzed the RNA-seq data. P.S. and B.N. provided the pCAP-SP1 peptide. M.E. and L.K.M provided the dnRARα plasmid. A.S.C and M.S. provided *GzmA/B* DKO samples and intellectual input. N.C., N.F. and E.dB. provided reagents and critical glycobiology input. L.C. and S.B. wrote the paper, with input from all the authors.

