## supplementary data for "A Glycosylation-Dependent Checkpoint Restrains Intestinal Intra-Epithelial Lymphocyte Activation"

Figure S1 *(Data from Nie et al., 2022)*

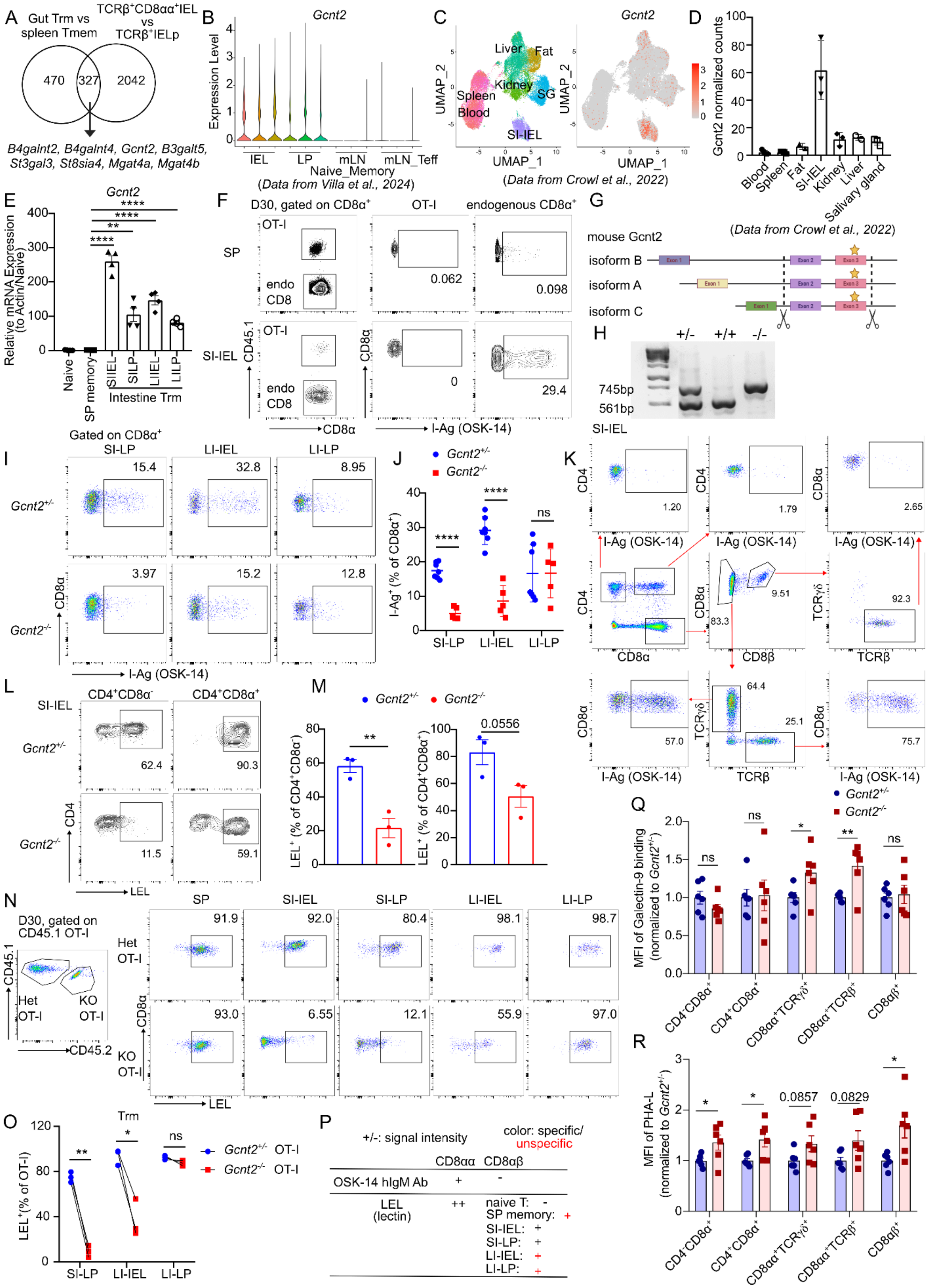

**Figure S1. GCNT2-meditated I-branching glycosylation is a defining feature of gut-resident $CD8^+$ T cells.**

(A) Number of upregulated genes from the analysis shown in Fig. 1C and Fig. 1D, depicting overlap between up-regulated genes in Trm OT-I cells and endogenous $TCR\beta^+CD8\alpha\alpha^+$ IEL cells.

(B) Reanalysis of *Gcnt2* expression across distinct $CD8^+$ T cell subsets from a published single-cell RNA-seq dataset (Villa et al., 2024).

(C) Reanalysis of *Gcnt2* expression in Trm cells isolated from different organs using a published scRNA-seq dataset combining P14 $CD8^+$ T cell transfer and LCMV infection (Crowl et al., 2022).

(D) Normalized counts of *Gcnt2* in Trm cells across distinct organs based on a published bulk RNA-seq dataset (Crowl et al., 2022).

(E) RT-qPCR result of *Gcnt2* expression in naïve, spleen memory and Trm OT-I cells from different intestinal compartments.

(F) Flow cytometry analysis of OSK-14 staining on memory OT-I cells and endogenous $CD8^+$ T cells from spleen and SI-IEL.

(G) Schematic representation of *Gcnt2* isoforms.

(H) Genotyping result of *Gcnt2*$^{-/-}$ mice generated by Crispr/Cas9.

(I) Representative FACS plot showing I-branching glycosylation detected by OSK-14 antibody staining on $CD8\alpha^+$ T cells from distinct intestinal compartments.

(J) Percentages of I-$Ag^+$ $CD8\alpha^+$ cells were quantified from (I).

(K) FACS plot showing OSK-14 staining of different endogenous SI-IEL subsets.

(L)-(M) FACS plot (L) and geometric mean fluorescence intensity (MFI) (M) of LEL staining on $CD4^+CD8\alpha\alpha^-$ and $CD4^+CD8\alpha\alpha^+$ SI-IEL.

(N) FACS plot showing LEL staining of splenic memory and Trm OT-I cells from different intestinal compartments at day 30 post infection.

(O) Quantification of $LEL^+$ OT-I cells from (N).

(P) Summary of staining intensity and specificity by using OSK-14 Ab and LEL.

(Q) MFI of Galectin-9 binding signal in each subset of SI-IEL, normalized to *Gcnt2*$^{+/-}$ group.

(R) MFI of lectin PHA-L binding signal in each subset of SI-IEL, normalized to *Gcnt2*$^{+/-}$ group.

Data in (E), (J), (Q) and (R) are pooled from two independent experiments with n=4 or n=5-6. Data in (M) are from one representative experiment of two independent experiments with n=3. Data in (O) are from one representative experiment of four independent experiments with n=3. Data are shown as mean ± SEM. Statistical analysis: unpaired t test. *p<0.05; **p<0.01; ****p<0.0001; ns, not significant.

FigureS2

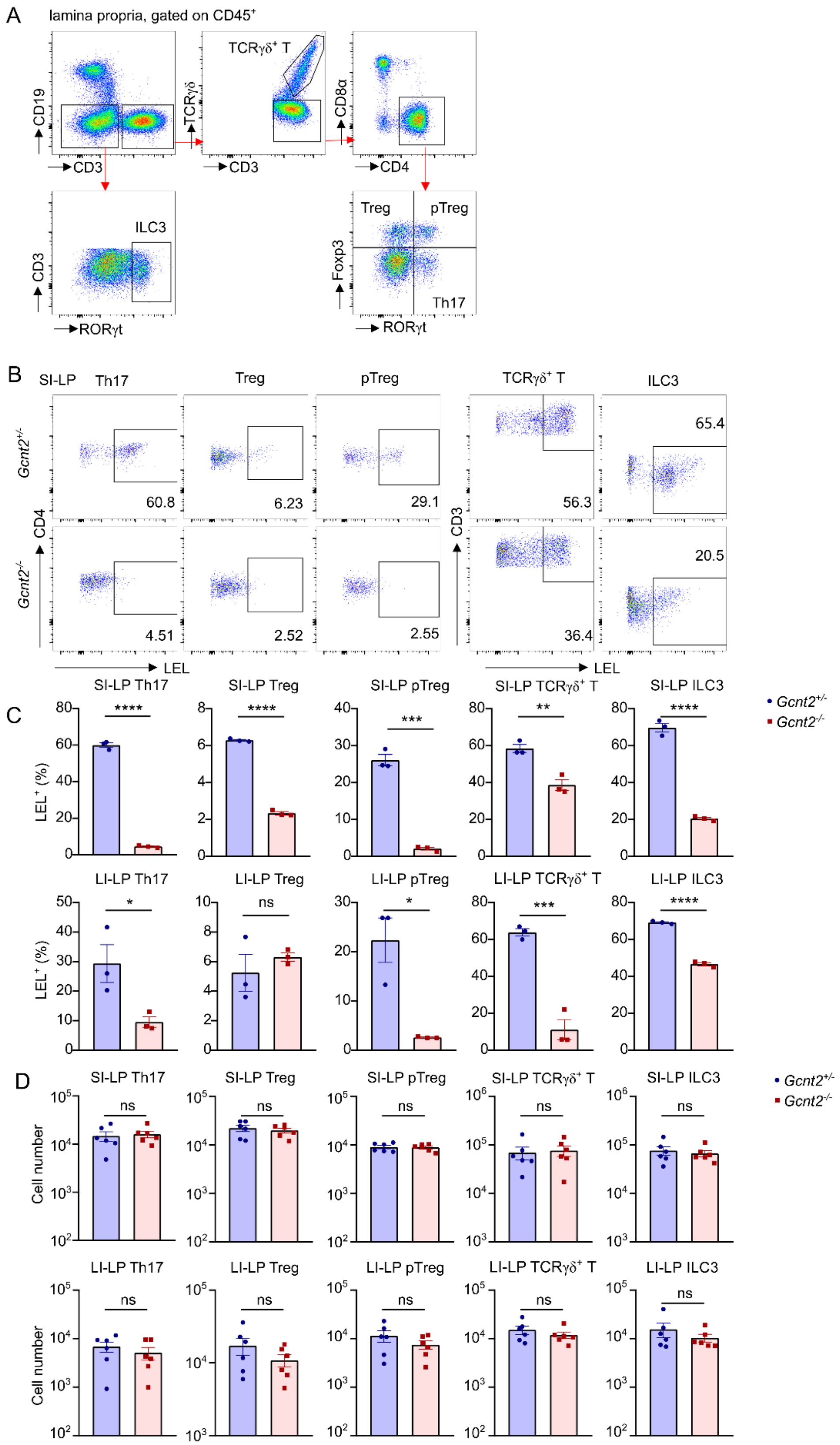
A
lamina propria, gated on CD45+
CD19
CD3
TCRγδ+ T
TCRγδ
CD8α
CD4
ILC3
RORγt
Treg
pTreg
Foxp3
Th17
B
SI-LP
Th17
Treg
pTreg
TCRγδ+ T
ILC3
Gcnt2+/-
Gcnt2-/-
CD4
CD3
LEL
60.8
6.23
29.1
56.3
65.4
4.51
2.52
2.55
36.4
20.5
C
SI-LP Th17
SI-LP Treg
SI-LP pTreg
SI-LP TCRγδ+ T
SI-LP ILC3
LI-LP Th17
LI-LP Treg
LI-LP pTreg
LI-LP TCRγδ+ T
LI-LP ILC3
LEL+ (%)
Gcnt2+/-
Gcnt2-/-
****
***
**
*
ns
D
Cell number
ns

**Figure S2. Display of GCNT2-dependent glycosylation on additional intestinal lymphoid populations.**

(A) Gating strategy for cells from small and large intestine lamina propria.

(B)-(C) FACS plots (B) and percentages (C) of LEL staining on distinct subsets in both SI and LI-LP.

(D) Quantification of cell number on distinct subsets in both SI and LI-LP.

Data in (C) is shown as one representative of two independent experiments with n=3. Data in (D) is pooled from two independent experiments with n=6. Data are shown as mean ± SEM. Statistical analysis: unpaired t test. *$p<0.05$; **$p<0.01$; ***$p<0.001$; ****$p<0.0001$; ns, not significant.

### Figure S3

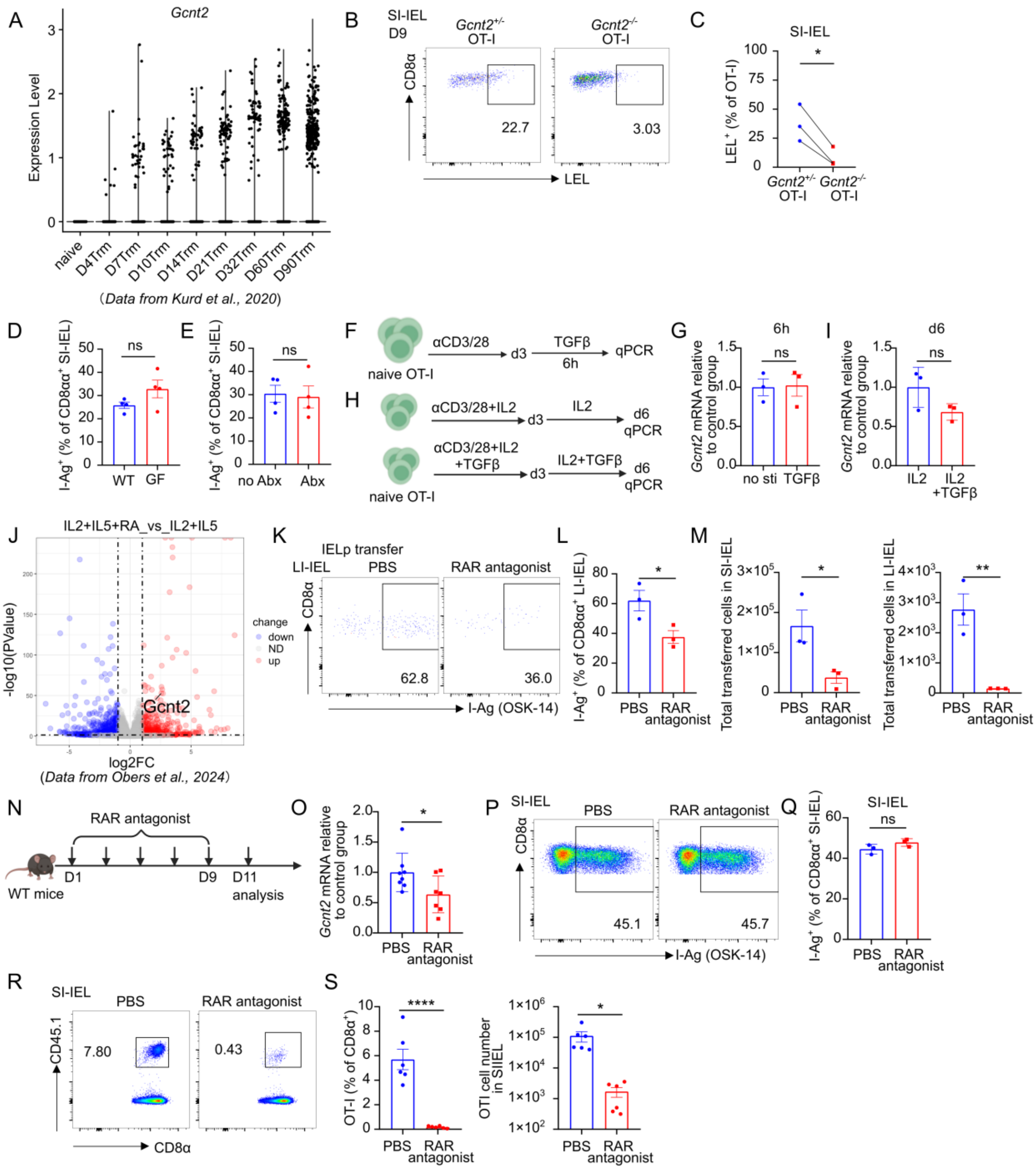
A
Gcnt2
Expression Level
naive
D4Trm
D7Trm
D10Trm
D14Trm
D21Trm
D32Trm
D60Trm
D90Trm
(Data from Kurd et al., 2020)
B
SI-IEL
D9
Gcnt2+/- OT-I
Gcnt2-/- OT-I
CD8α
LEL
22.7
3.03
C
SI-IEL
LEL+ (% of OT-I)
Gcnt2+/- OT-I
Gcnt2-/- OT-I
D
I-Ag+ (% of CD8αα+ SI-IEL)
ns
WT
GF
E
I-Ag+ (% of CD8αα+ SI-IEL)
ns
no Abx
Abx
F
naive OT-I
αCD3/28
d3
TGFβ
6h
qPCR
H
αCD3/28+IL2
d3
IL2
d6
qPCR
αCD3/28+IL2
+TGFβ
d3
IL2+TGFβ
d6
qPCR
naive OT-I
G
6h
Gcnt2 mRNA relative to control group
ns
no sti
TGFβ
I
d6
Gcnt2 mRNA relative to control group
ns
IL2
IL2
+TGFβ
J
IL2+IL5+RA_vs_IL2+IL5
-log10(PValue)
change
down
ND
up
Gcnt2
log2FC
(Data from Obers et al., 2024)
K
IELp transfer
LI-IEL
PBS
RAR antagonist
CD8α
62.8
36.0
I-Ag (OSK-14)
L
I-Ag+ (% of CD8αα+ LI-IEL)
PBS
RAR antagonist
M
Total transferred cells in SI-IEL
PBS
RAR antagonist
Total transferred cells in LI-IEL
PBS
RAR antagonist
N
RAR antagonist
WT mice
D1
D9
D11
analysis
O
Gcnt2 mRNA relative to control group
PBS
RAR antagonist
P
SI-IEL
PBS
RAR antagonist
CD8α
45.1
45.7
I-Ag (OSK-14)
Q
SI-IEL
ns
I-Ag+ (% of CD8αα+ SI-IEL)
PBS
RAR antagonist
R
SI-IEL
PBS
RAR antagonist
CD45.1
7.80
0.43
CD8α
S
OT-I (% of CD8α+)
PBS
RAR antagonist
OTI cell number in SIIEL
PBS
RAR antagonist

**Figure S3. Retinoic acid promotes *Gcnt2* expression.**

(A) *Gcnt2* expression in SI-IEL Trm cells at different time points from a published scRNA-seq dataset utilizing P14 cell transfer and LCMV infection (Kurd et al., 2020).

(B-C) CD45.1 *Gcnt2*$^{+/-}$ and CD45.1/2 *Gcnt2*$^{-/-}$ OT-I cells were co-transferred into CD45.2 WT recipients, followed by oral *Lm-OVA* infection. At day 9, LEL staining on SI-IEL OT-I cells was analyzed by FACS (B) and quantified (C).

(D) Percentage of I-Ag$^+$ SI-IEL CD4$^-$CD8αα$^+$ T cells in WT specific pathogen free (SPF) and germ-free (GF) mice.

(E) WT mice were administered a cocktail of broad-spectrum antibiotics (ampicillin, metronidazole, neomycin and vancomycin) in drinking water for one month. Percentage of I-Ag$^+$ SI-IEL CD4$^-$CD8αα$^+$ T cells were quantified.

(F) Enriched splenic OT-I cells were activated with plate-bound anti-CD3/28 for three days, then stimulated with TGF-β (10 ng/ml) for six hours.

(G) RT-qPCR was performed to detect *Gcnt2* expression as described in (F).

(H) For long-term stimulation, cells were cultured with TGF-β for six days.

(I) RT-qPCR was performed to detect *Gcnt2* expression as described in (H).

(J) Reanalysis of *Gcnt2* expression in RA-treated cells from a published bulk RNA-seq dataset (Obers et al., 2024).

(K)-(L) I-branching glycosylation staining using OSK-14 antibody on LI-IEL CD8αα$^+$ T cells was detected by FACS (K) from vehicle or RAR antagonist treated *Rag2*$^{-/-}$ mice receiving WT IEL precursors transfer, and percentage of I-Ag$^+$ LI-IEL CD8αα$^+$ T cells were quantified (L).

(M) Total number of transferred cells in SI-IEL and LI-IEL were quantified, referring to Fig. 2K.

(N) Experimental layout. WT mice receiving vehicle or RAR antagonist treatment (1 mg/kg) every other day for 7 days.

(O) SI-IEL CD8αα$^+$ T cells were sorted from (N) and RT-qPCR was performed to detect *Gcnt2* expression level.

(P)-(Q) I-branching glycosylation staining using OSK-14 antibody on SI-IEL CD8αα$^+$ T cells was detected by FACS (P) and quantified (Q), as described in (N).

(R-S) In the CD45.1 OT-I transfer model with RAR antagonist treatment, SI-IEL OT-I cells were analyzed by FACS (R) and quantified (S), referring to Fig. 2N.

Data in (C), (D), (E), (G), (I), (L), (M) and (Q) represent one of two independent experiments with n=3-4. Data in (O) are pooled from three independent experiments. Data in (S) are pooled from two independent experiments. Data are shown as mean ± SEM. Statistical analysis: unpaired t test. *p<0.05; **p<0.01; ****p<0.0001; ns, not significant.

Figure S4

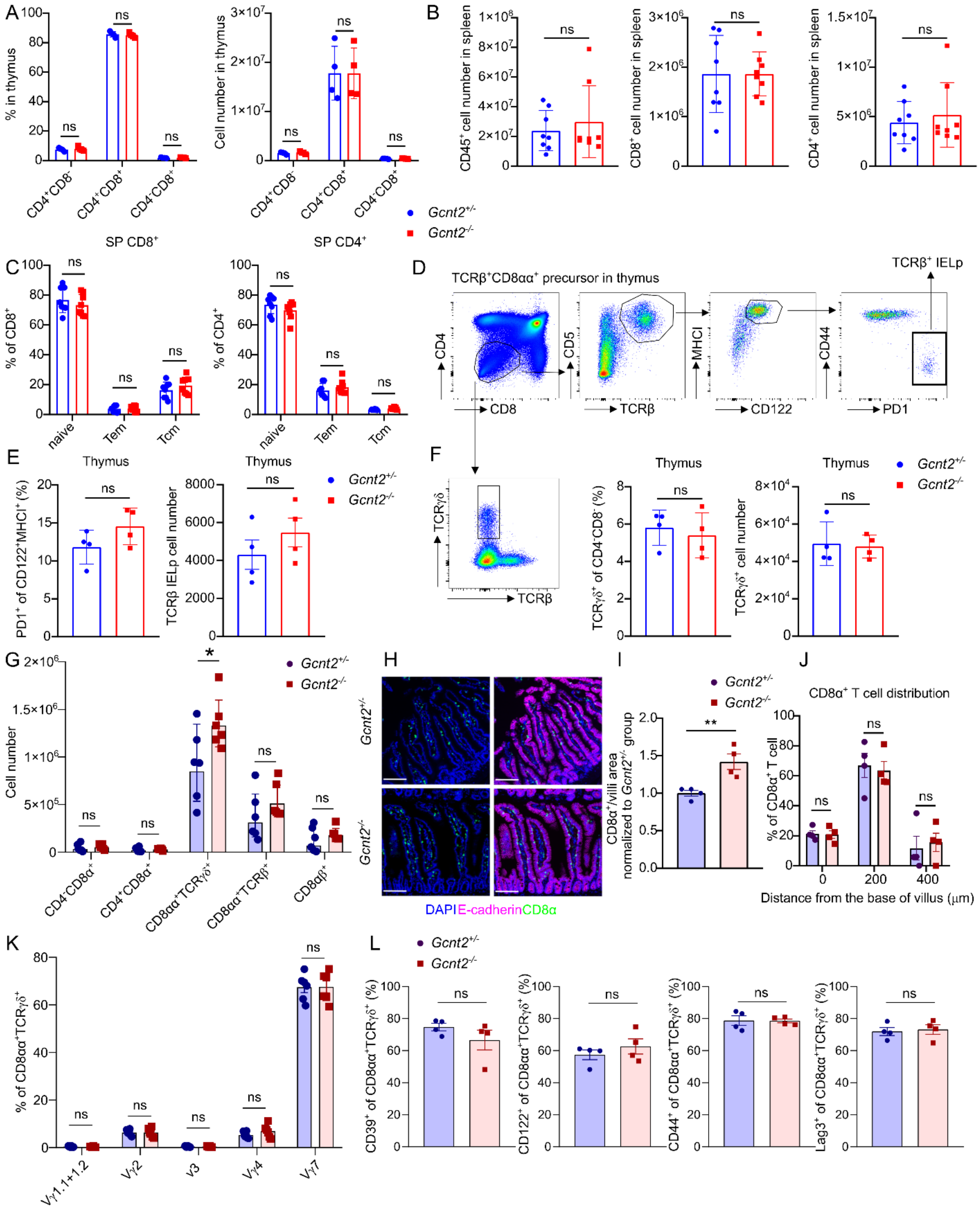


100

**Figure S4. Thymic and peripheral T cell compartments analysis in *Gcnt2*$^{-/-}$ mice.**

(A) Percentages and numbers of CD4$^{+}$CD8$^{-}$, CD4$^{+}$CD8$^{+}$ and CD4$^{-}$CD8$^{+}$ T cells in the thymus of 6-8-week-old *Gcnt2*$^{+/-}$ and *Gcnt2*$^{-/-}$ mice.

(B) Total numbers of CD45$^{+}$, CD4$^{+}$ and CD8$^{+}$ cells in the spleen of 6-8-week-old *Gcnt2*$^{+/-}$ and *Gcnt2*$^{-/-}$ mice.

(C) Percentage of naïve (CD44$^{-}$CD62L$^{+}$), effector memory (CD44$^{+}$CD62L$^{-}$) and central memory (CD44$^{-}$CD62L$^{+}$) subsets among CD4$^{+}$ and CD8$^{+}$ T cells in the spleen.

(D) Gating strategy to identify CD8αα$^{+}$TCRβ$^{+}$ IEL precursors in the thymus.

(E) Percentages and numbers of CD8αα$^{+}$TCRβ$^{+}$ IEL precursors in the thymus of 6-8-week-old *Gcnt2*$^{+/-}$ and *Gcnt2*$^{-/-}$ mice.

(F) FACS plot showing TCRγδ staining within the CD4$^{-}$CD8$^{-}$ subset in the thymus (left). Percentages and numbers of TCRγδ$^{+}$ IEL precursor cells in the thymus of 6-8-week-old *Gcnt2*$^{+/-}$ and *Gcnt2*$^{-/-}$ mice.

(G) Number of different subset in SI-IEL of 6-8-week-old *Gcnt2*$^{+/-}$ and *Gcnt2*$^{-/-}$ mice.

(H) Immunofluorescence staining of CD8α in the small intestine.

(I) Normalized number of CD8α$^{+}$ T cell per villus area (See Methods).

(J) Quantification of CD8α$^{+}$ T cell distance from the villus base.

(K) Vγ usage of CD4$^{-}$CD8αα$^{+}$TCRγδ$^{+}$ SI-IEL.

(L) Activation marker expression on CD4$^{-}$CD8αα$^{+}$TCRγδ$^{+}$ SI-IEL from *Gcnt2*$^{+/-}$ and *Gcnt2*$^{-/-}$ mice.

Data in (A), (E), (F), (G), (K) and (L) are pooled from two independent experiments with n=4 or n=6. Data in (B) and (C) are pooled from three independent experiments with n=8. Each dot in (I) and (J) represents one mouse, with values averaged from 10 villi. Data are shown as mean ± SEM. Statistical analysis: unpaired t test. *p<0.05; **p<0.01; ns, not significant.

Figure S5

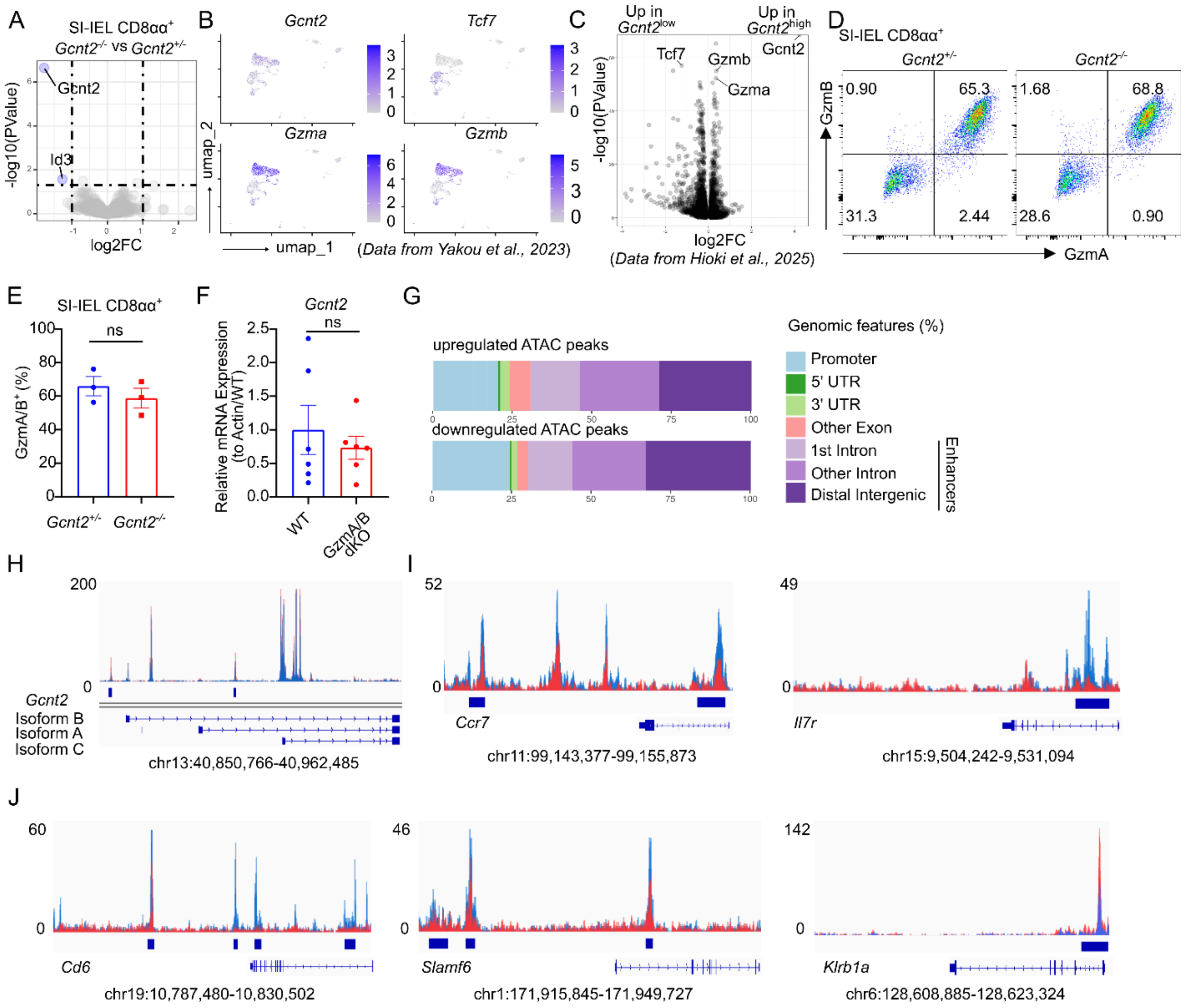


133

**Figure S5. Distinct chromatin accessibility profiles in I-Ag$^+$ and I-Ag$^-$ CD8αα$^+$ SI-IELs.**

(A) Volcano plot depicting differentially expressed genes in SI-IEL CD4$^-$CD8αα$^+$ T cells between *Gcnt2*$^{+/-}$ and *Gcnt2*$^{-/-}$ mice.

(B) Reanalysis of IEL scRNA-seq dataset showing *Gcnt2*, *Tcf7*, *Gzma* and *Gzmb* expression (Yakou et al., 2023).

(C) Reanalysis of published IEL scRNA-seq dataset showing correlated expression of *Gcnt2* with *Gzma* and *Gzmb* (Hioki et al., 2025).

(D) FACS plot of GzmA/B expression in CD4$^-$CD8αα$^+$ SI-IEL from *Gcnt2*$^{+/-}$ and *Gcnt2*$^{-/-}$ mice.

(E) Quantification of data shown in (D).

(F) RT-qPCR analysis of *Gcnt2* expression in sorted CD4$^-$CD8αα$^+$ SI-IEL from WT and *Gzma/b* double knockout (dKO) mice.

(G) Genomic distribution of differentially accessible peaks identified by ATAC-seq.

(H) IGV browser view of chromatin accessibility at *Gcnt2* locus.

(I) IGV browser view of chromatin accessibility at the *Ccr7* and *Il7r* loci.

(J) IGV browser view of chromatin accessibility at the loci for co-stimulatory molecules *Cd6* and *Slamf6*, as well as inhibitory molecule *Klrb1a*.

Data in (E) is one representative of two independent experiments with n=3. Data in (F) is pooled from six mice per group. Data are shown as mean ± SEM. Statistical analysis: unpaired t test. ns, not significant.

Figure S6

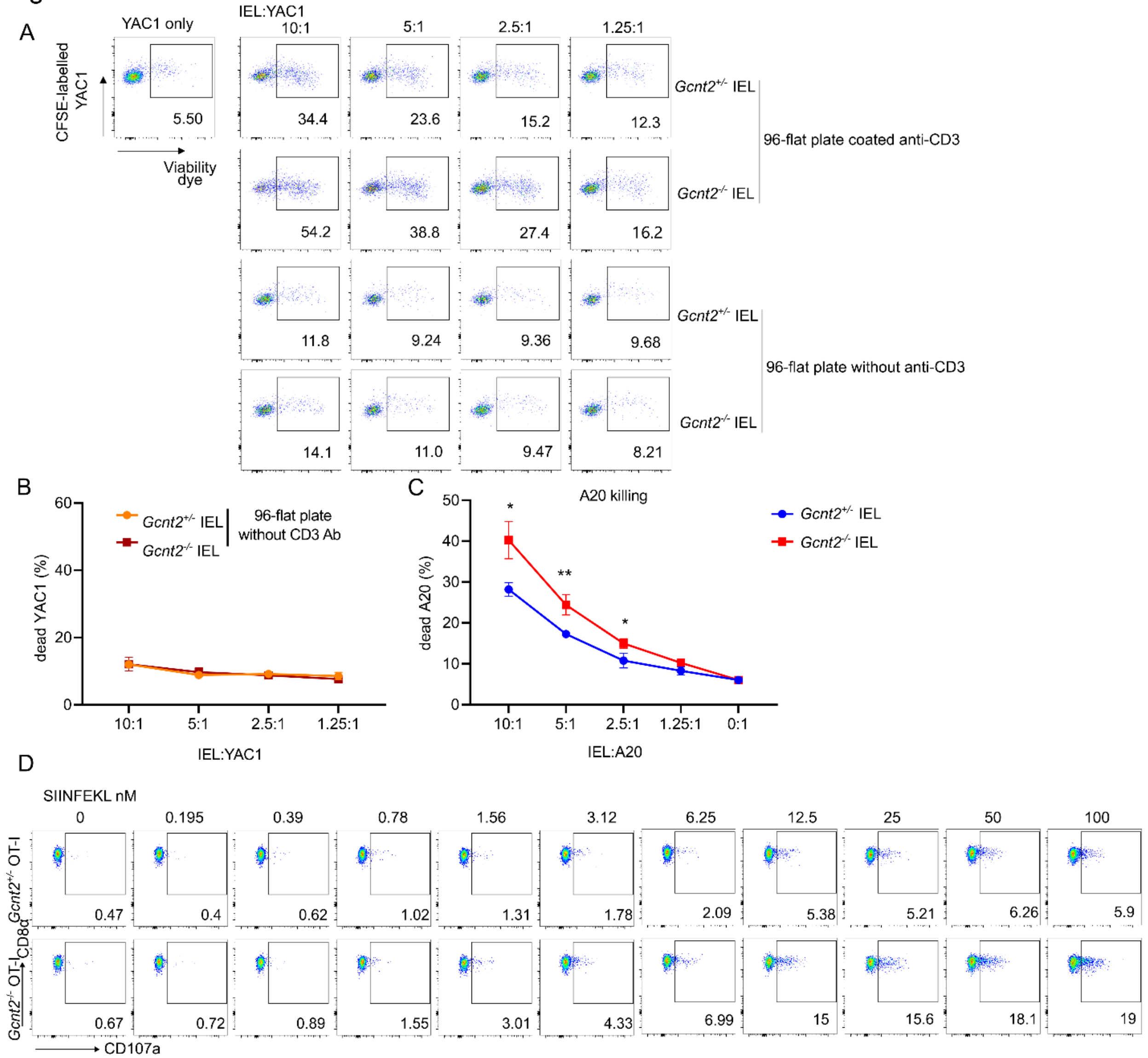


158

**Figure S6. *Gcnt2* deficient CD8αα$^+$ SI-IELs display enhanced cytotoxicity.**

(A) FACS plot showing dead YAC1 cells after co-culture with SI-IEL CD4$^-$CD8αα$^+$ T cells in the presence or absence of anti-CD3 stimulation.

(B) Percentage of dead YAC1 target cells after co-culture with non-activated CD4$^-$CD8αα$^+$ IELs at the indicated effector-to-target ratios.

(C) Percentage of dead A20 target cells after co-culture with anti-CD3-activated CD4$^-$CD8αα$^+$ IELs at the indicated effector-to-target ratios.

(D) Representative FACS plot showing degranulation of SI-IEL CD8αβ$^+$ T cells from OT-I background mice following SIINFEKL stimulation, referring to Fig. 4J.

Data in (B) and (C) were one representative of two independent experiments. Data are shown as mean ± SEM. Statistical analysis: unpaired t test. *$p<0.05$; **$p<0.01$.

Figure S7

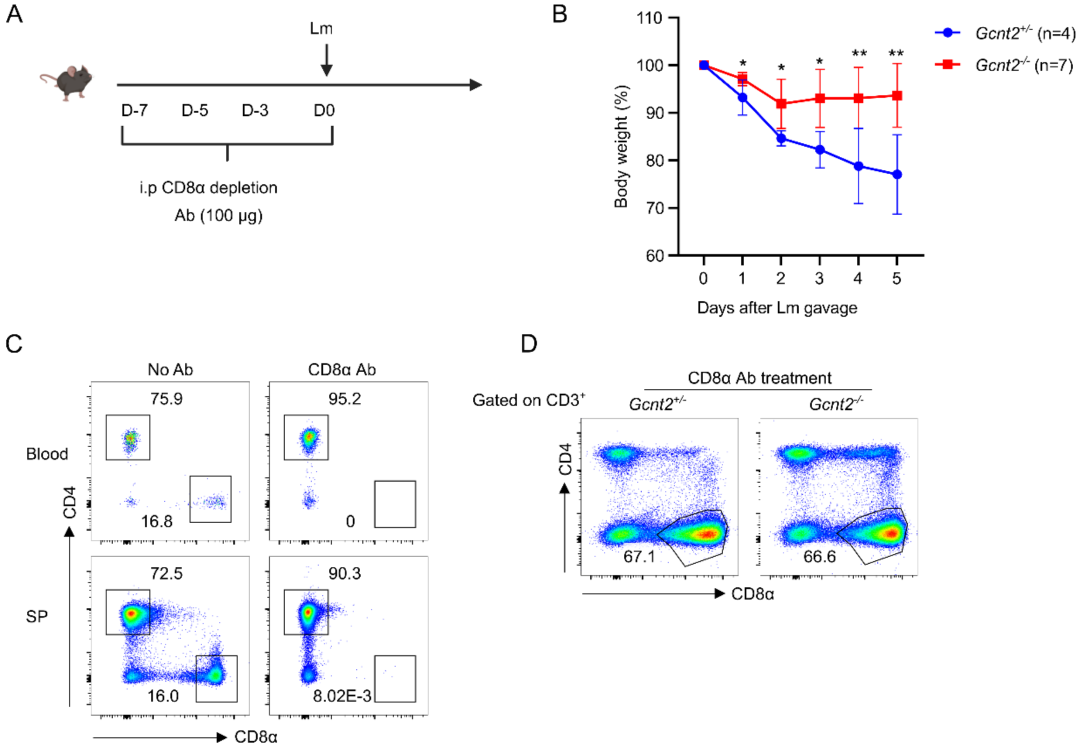


186

**Figure S7. Depletion of circulating CD8$^+$ T cells fails to affect *Gcnt2*-dependent phenotype.**

(A) *Gcnt2*$^{+/-}$ or *Gcnt2*$^{-/-}$ mice were administrated with CD8α depletion antibody (100 µg) via intraperitoneal injection from day -7 to day 0, followed by *Listeria monocytogenes* (1x10$^8$) oral infection.

(B) Body weight change in mice following *Listeria monocytogenes* oral infection, as described in (A).

(C-D) Depletion efficiency of CD8α$^+$ T cells was assessed by flow cytometry in peripheral blood and spleen (C) and in the small intestine (D) at day 0.

Data in (B-D) were one representative of two independent experiments. Data are shown as mean ± SEM. Statistical analysis: unpaired t test. *p<0.05; **p<0.01; ns, not significant.

Figure S8

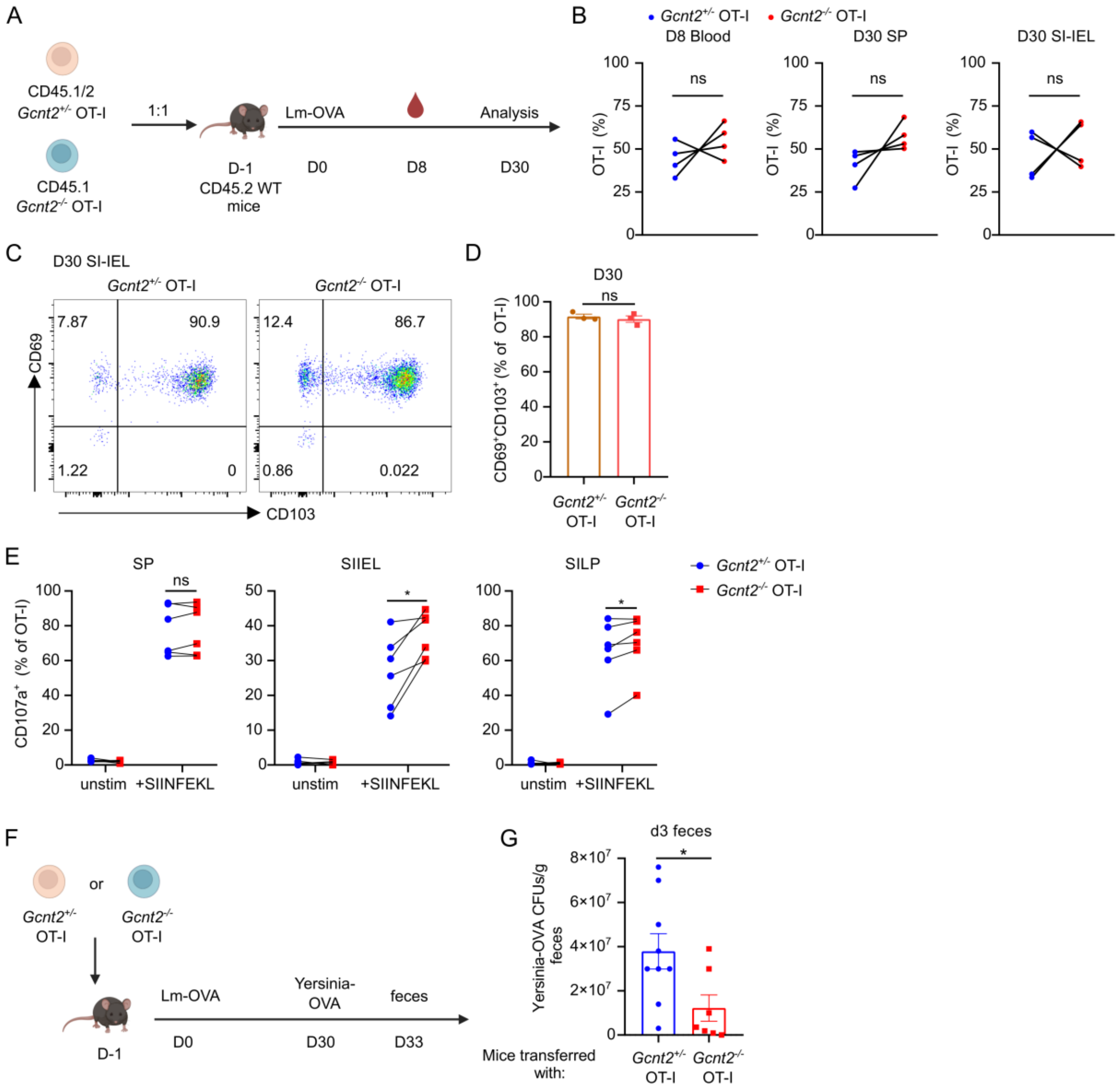
A
CD45.1/2
Gcnt2+/- OT-I
CD45.1
Gcnt2-/- OT-I
1:1
D-1
CD45.2 WT
mice
Lm-OVA
D0
D8
Analysis
D30
B
Gcnt2+/- OT-I
Gcnt2-/- OT-I
D8 Blood
D30 SP
D30 SI-IEL
OT-I (%)
ns
C
D30 SI-IEL
Gcnt2+/- OT-I
Gcnt2-/- OT-I
7.87
90.9
1.22
0
12.4
86.7
0.86
0.022
CD69
CD103
D
D30
ns
CD69+CD103+ (% of OT-I)
E
SP
SIIEL
SILP
CD107a+ (% of OT-I)
unstim
+SIINFEKL
F
or
Gcnt2+/- OT-I
Gcnt2-/- OT-I
D-1
Lm-OVA
D0
Yersinia-OVA
D30
feces
D33
G
d3 feces
Yersinia-OVA CFUs/g feces
Mice transferred with:

**Figure S8. *Gcnt2* deficient Trm cells provide superior protection against reinfection.**

(A) *Gcnt2*$^{+/-}$ and *Gcnt2*$^{-/-}$ OT-I cells with different congenic markers were co-transferred into CD45.2 WT recipients at a 1:1 ratio, followed by oral infection with *Lm-OVA*.

(B) The ratio of donor cells in peripheral blood was analyzed at day 8, and in the spleen and small intestine at day 30 post infection by FACS.

(C)-(D) Expression of CD69 and CD103 on co-transferred OT-I cells at day 30 (C) and corresponding quantification (D).

(E) At day 30, cells from the indicated organs in (A) were isolated and co-cultured with SIINFEKL-loaded CD45.2 splenocytes in the presence of CD107a antibody and Golgi inhibitor for 4 h. Degranulation, indicated by CD107a surface expression, was analyzed by FACS.

(F)-(G) *Gcnt2*$^{+/-}$ and *Gcnt2*$^{-/-}$ OT-I cells were transferred to WT recipients separately, followed by oral *Listeria-OVA* infection. At day 30, mice were secondarily infected orally with *Yersinia-OVA* ($3x10^8$). Fecal samples were collected at day 3 post reinfection and plated on *Yersinia*-selective agar plates for bacterial load quantification (G).

Data in (B) are pooled from four independent experiments. Data in (D) represent one of four independent experiments with n=3. Data in (E) are pooled from two independent experiments with n=6. Data in (G) are pooled from two independent experiments with n=7-9. Data are shown as mean ± SEM. Statistical analysis: unpaired t test. *$p<0.05$; ns, not significant.

Figure S9

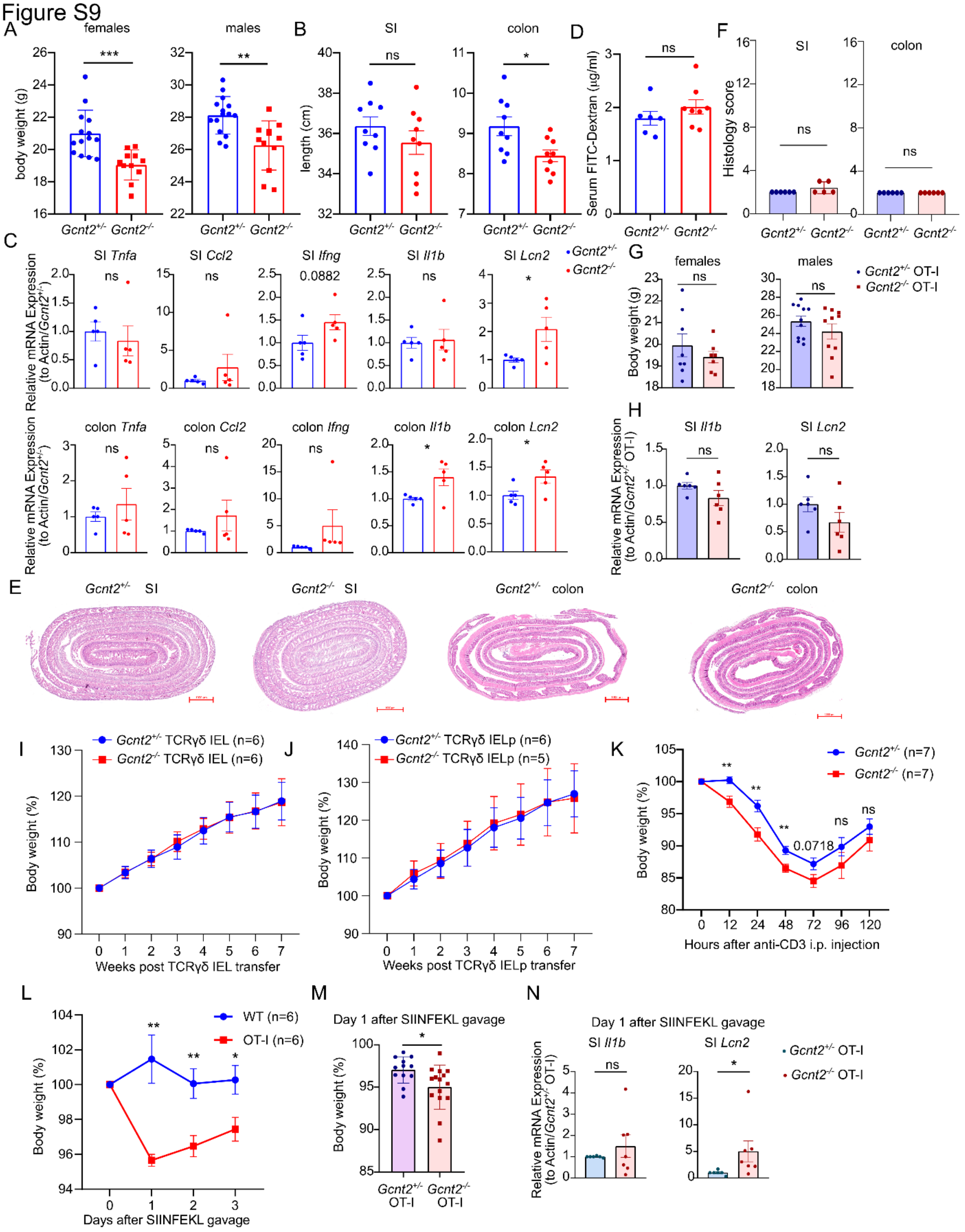


249

**Figure S9. *Gcnt2* deficiency exhacerbates TCR activation-induced inflammation.**

(A) Body weight of male and female *Gcnt2*$^{+/-}$ and *Gcnt2*$^{-/-}$ mice at 10 weeks of age.

(B) Small intestine and colon length in *Gcnt2*$^{+/-}$ and *Gcnt2*$^{-/-}$ mice at 10 weeks of age.

(C) RT-qPCR of inflammatory genes in small intestine or colon from 10-week old *Gcnt2*$^{+/-}$ and *Gcnt2*$^{-/-}$ mice at steady state.

(D) Serum 4-kDa FITC dextran levels in *Gcnt2*$^{+/-}$ and *Gcnt2*$^{-/-}$ mice, indicating intestinal barrier permeability.

(E) HE staining of small intestine (duodenum) and colon from *Gcnt2*$^{+/-}$ and *Gcnt2*$^{-/-}$ mice. Scale bar represents 100 µm.

(F) Histology score of HE staining results from (E).

(G) Body weight of both 10-week old male and female *Gcnt2*$^{+/-}$ and *Gcnt2*$^{-/-}$ OT-I mice at steady state.

(H) RT-qPCR of inflammatory genes in small intestine from 10-week old *Gcnt2*$^{+/-}$ and *Gcnt2*$^{-/-}$ OT-I mice at steady state.

(I) A total of $5 \times 10^5$ $CD4^-CD8\alpha\alpha^+TCR\gamma\delta^+$ SI-IEL cells isolated from *Gcnt2*$^{+/-}$ or *Gcnt2*$^{-/-}$ mice were adoptively transferred into *Rag2*$^{-/-}$ recipient mice, and body weight was monitored thereafter.

(J) A total of $5 \times 10^4$ $CD4^-CD8^-TCR\gamma\delta^+$ thymocytes isolated from *Gcnt2*$^{+/-}$ or *Gcnt2*$^{-/-}$ mice were adoptively transferred into *Rag2*$^{-/-}$ recipient mice, and body weight was monitored thereafter.

(K) *Gcnt2*$^{+/-}$ and *Gcnt2*$^{-/-}$ mice were intraperitoneally injected with 25 µg anti-CD3 antibody. Body weight changes were monitored over time.

(L) WT and OT-I mice were orally gavaged with SIINFEKL peptide (100 µg). Body weight change was monitored.

(M) Body weight change of *Gcnt2*$^{+/-}$ or *Gcnt2*$^{-/-}$ OT-I mice at day one post SIINFEKL peptide gavage.

(N) RT-qPCR analysis of inflammatory gene expression in small intestine at day one post oral gavage of SIINFEKL peptide.

Data in (A) are pooled from 11-15 mice per group. Data in (B) are pooled from 9 mice per group. Data in (C) are one representative of two independent experiments. Data in (E) are pooled from 5-6 mice. Data in (F) is one representative of two independent experiments. Data in (G) is pooled from 7-11 mice. Data in (H), (I), (J), (K), (L) and (N) are pooled from 5-7 mice per group. Data in (M) is pooled from 12-15 mice per group. Data are shown as mean ± SEM. Statistical analysis: unpaired t test. *$p<0.05$; **$p<0.01$; ***$p<0.001$; ns, not significant.

Figure S10

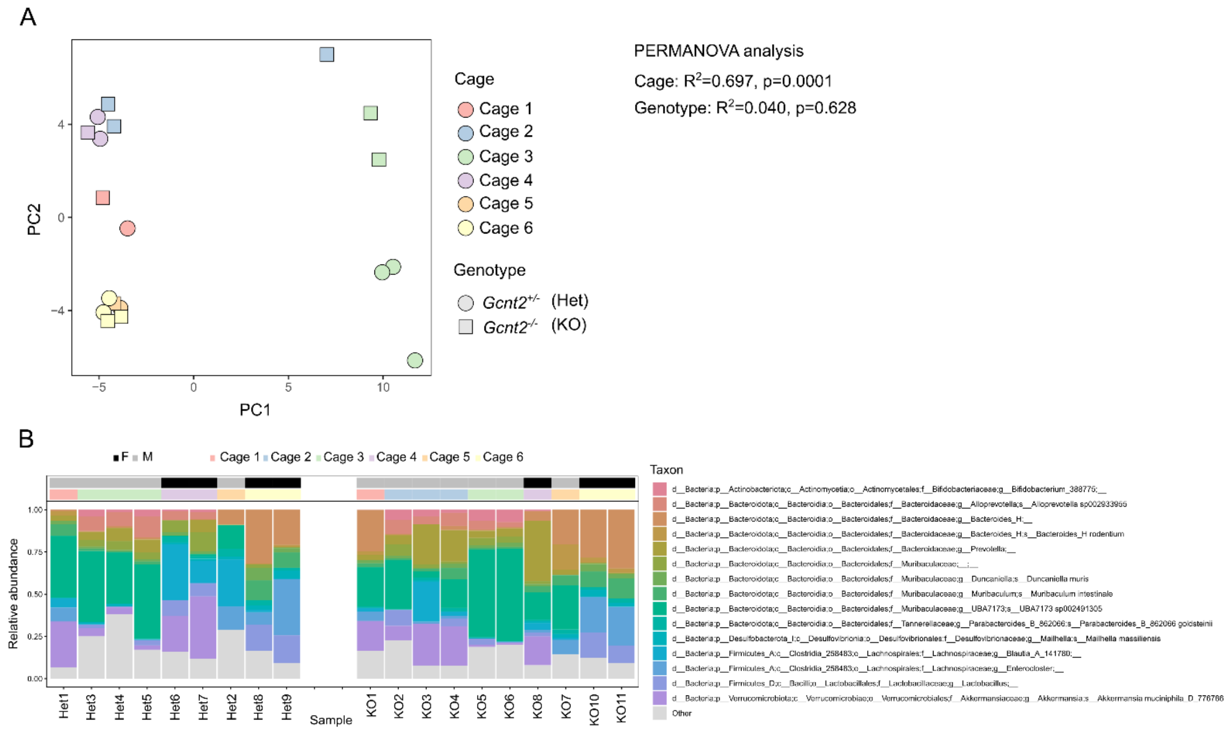


282

**Figure S10. 16S rRNA sequencing analysis of fecal microbiota.**

(A) 16S rRNA gene profiling did not reveal a major genotype-associated shift in fecal microbiota composition, as genotype explained only a minor and non-significant fraction of heta-diversity variation both alone (PERMANOVA, $R^2$=0.040, p=0.628) and after accounting for cage and other available covariates (marginal PERMANOVA, $R^2$=0.030, p=0.221), whereas cage-associated variation dominated the dataset ($R^2$=0.697, p=0.0001).

(B) Taxonomic profiles of the fecal microbiota based on 16S rRNA sequencing of samples from *Gcnt2*$^{+/-}$ and *Gcnt2*$^{-/-}$ mice.

Figure S11

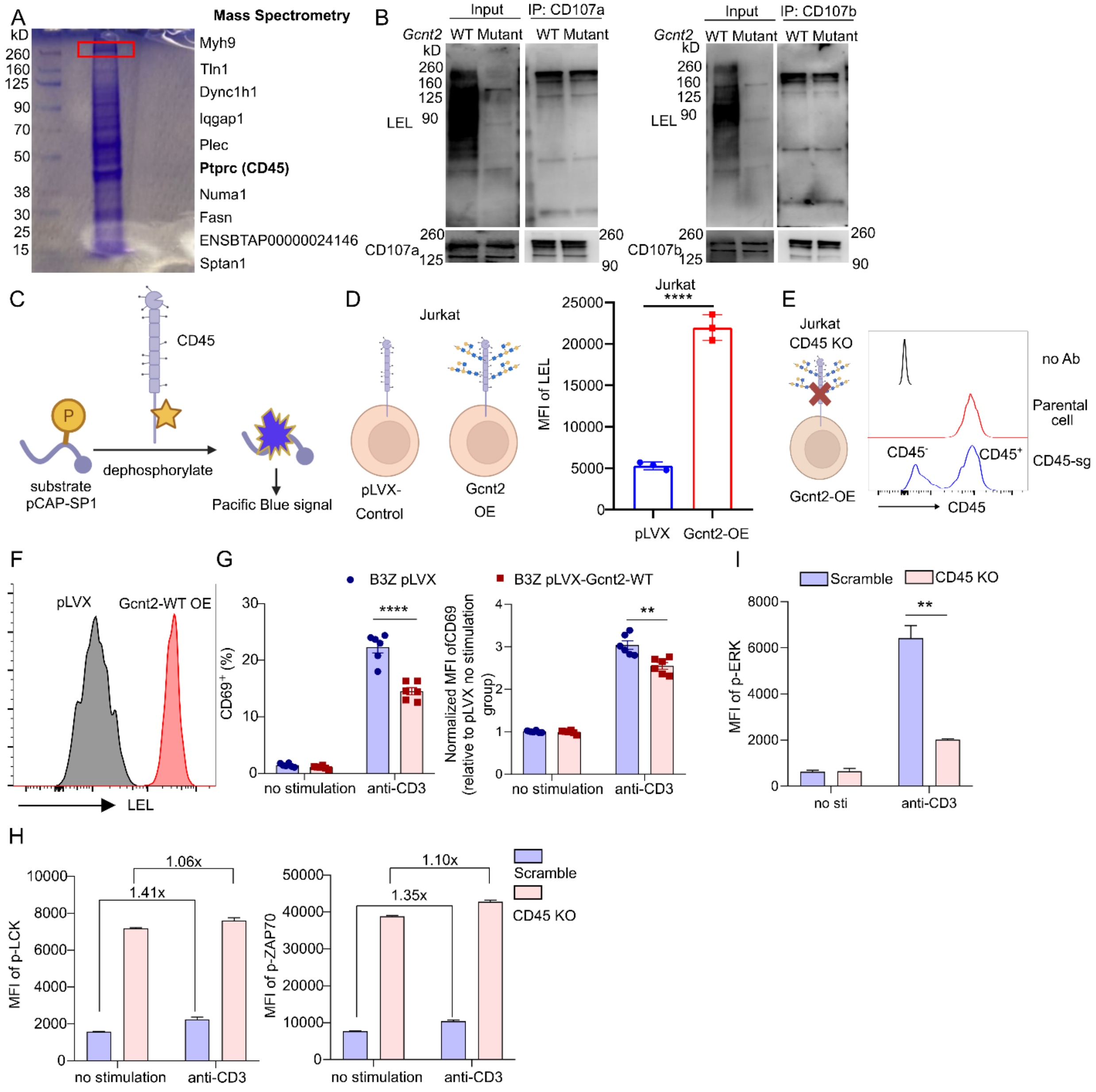
A
Mass Spectrometry
kD
260
160
125
90
70
50
38
30
25
15
Myh9
Tln1
Dync1h1
Iqgap1
Plec
Ptprc (CD45)
Numa1
Fasn
ENSBTAP00000024146
Sptan1
B
Input
IP: CD107a
Gcnt2
WT Mutant
WT Mutant
LEL
CD107a
Input
IP: CD107b
CD107b
C
CD45
P
substrate pCAP-SP1
dephosphorylate
Pacific Blue signal
D
Jurkat
pLVX-Control
Gcnt2 OE
MFI of LEL
pLVX
Gcnt2-OE
****
E
Jurkat CD45 KO
Gcnt2-OE
no Ab
Parental cell
CD45-sg
CD45⁻
CD45⁺
CD45
F
pLVX
Gcnt2-WT OE
LEL
G
B3Z pLVX
B3Z pLVX-Gcnt2-WT
CD69⁺ (%)
Normalized MFI ofCD69 (relative to pLVX no stimulation group)
no stimulation
anti-CD3
****
**
I
Scramble
CD45 KO
MFI of p-ERK
no sti
anti-CD3
**
H
MFI of p-LCK
MFI of p-ZAP70
1.06x
1.41x
1.10x
1.35x
Scramble
CD45 KO
no stimulation
anti-CD3

**Figure S11. GCNT2 glycosylates CD45 to restrain TCR signaling.**

(A) Coomassie blue staining of *Gcnt2*$^{+/-}$ total SI-IEL CD8α$^{+}$ T cell lysates. The indicated gel band was excised and subjected to mass spectrometry to identify potential targets. Depicted are the top 10 hits identified by this analysis.

(B) CD107a (left) and CD107b (right) were immunoprecipitated from *Gcnt2*$^{-/-}$ OT-I cell lysates overexpressing *Gcnt2* WT or mutant forms, followed by LEL blotting.

(C) Schematic representation of the assay to detect CD45 phosphatase activity using pCAP-SP1 probe.

(D) Schematic representation of *Gcnt2* overexpression in Jurkat cell line and MFI of LEL staining on Jurkat cells transfected with empty vector or *Gcnt2* construct.

(E) Crispr/Cas9-mediated CD45 knockout in *Gcnt2*-overexpressing Jurkat cells.

(F) MFI of LEL staining on B3Z cells transfected with empty vector or *Gcnt2* construct.

(G) CD69 percentage (left) and MFI (right) 3 h after plate-coated anti-CD3 (5 µg/ml) stimulation in B3Z cells expressing empty vector or *Gcnt2*.

(H) MFI of phosphorylated LCK (Y394) and ZAP70 (Y319) at baseline or 10 min after anti-CD3 (5 µg/ml) stimulation in control and CD45 KO Jurkat cells.

(I) MFI of phosphorylated ERK (T202/Y204) at baseline or 10 min after anti-CD3 (5 µg/ml) stimulation in control and CD45 KO Jurkat cells.

Data in (B), (D), (H) and (I) were one representative of two independent experiments. Data in (G) was pooled from two independent experiments. Data are shown as mean ± SEM. Statistical analysis: unpaired t test for the data. $^{**}p<0.01$; $^{****}p<0.0001$.
